# The most widespread phage in animals: Genomics and taxonomic classification of Phage WO

**DOI:** 10.1101/2021.11.04.467296

**Authors:** Sarah R. Bordenstein, Seth R. Bordenstein

## Abstract

*Wolbachia* are the most common obligate, intracellular bacteria in animals. They exist worldwide in arthropod and nematode hosts in which they commonly act as reproductive parasites or mutualists, respectively. Bacteriophage WO, the largest of *Wolbachia*’s mobile elements, includes reproductive parasitism genes, serves as a hotspot for genetic divergence and genomic rearrangement of the bacterial chromosome, and uniquely encodes a Eukaryotic Association Module with eukaryotic-like genes and an ensemble of putative host interaction genes. Despite WO’s relevance to genome evolution, selfish genetics, and symbiotic applications, relatively little is known about its origin, host range, diversification, and taxonomic classification. Here we analyze the most comprehensive set of 150 *Wolbachia* and phage WO assemblies to provide a framework for discretely organizing and naming integrated phage WO genomes. We demonstrate that WO is principally in arthropod *Wolbachia* with relatives in diverse endosymbionts and metagenomes, organized into four variants related by gene synteny, often oriented opposite the origin of replication in the *Wolbachia* chromosome, and the large serine recombinase is an ideal typing tool to assign taxonomic classification of the four variants. We identify a novel, putative lytic cassette and WO’s association with a conserved eleven gene island, termed Undecim Cluster, that is enriched with virulence-like genes. Finally, we evaluate WO-like Islands in the *Wolbachia* genome and discuss a new model in which Octomom, a notable WO-like Island, arose from a split with WO. Together, these findings establish the first comprehensive Linnaean taxonomic classification of endosymbiont phages that includes distinguishable genera of phage WO, a family of non-*Wolbachia* phages from aquatic environments, and an order that captures the collective relatedness of these viruses.

## Introduction

Intracellular, endosymbiotic bacteria comprise some of the most intimate and enduring host-microbe interactions. While reductive evolutionary forces are often presumed to lead to streamlined, tiny genomes, many endosymbionts that host switch contain notable levels of active or relic mobile DNA [1]. An exemplar is the genus *Wolbachia* which harbor transposons [2], temperate phages [3, 4], and putative plasmids [5, 6]. *Wolbachia* are members of the Anaplasmataceae family [7] that also includes the intracellular genera *Anaplasma, Ehrlichia, Neorickettsia, Aegptianella*, and several newly classified bacteria. *Wolbachia* occur in a vast number of invertebrates spanning some nematodes and roughly half of all arthropod species, thus making them the most widespread endosymbionts in animals [8]; but unlike its sister genera, it does not naturally occur in mammalian hosts [9]. Transmission routes are predominantly vertical through the germline, and horizontal transmission of *Wolbachia* in arthropods is frequent on an evolutionary timescale [10, 11], leading to coinfections and subsequent bacteriophage exchanges in the same host [12–16]. Integrated within the bacterial chromosome, these bacteriophages are hot spots of genetic divergence between *Wolbachia* strains [6, 17–20].

Many arthropod-associated *Wolbachia* cause various forms of reproductive parasitism including feminization, parthenogenesis, male killing, and cytoplasmic incompatibility (CI). These selfish modifications hijack sex determination, sex ratios, gametogenesis, and/or embryonic viability to enhance the spread of *Wolbachia* through the transmitting matriline [21, 22]. Nematode-associated *Wolbachia*, however, generally lack phage WO and more often act as mutualists within their animal host [23, 24]. Thus, phage WO was originally hypothesized to contribute to these reproductive manipulations in arthropods through horizontal acquisition and differential expression of parasitism genes that are not part of the core *Wolbachia* genome [20, 23, 25–28]. Indeed, transgenic expression of two genes from phage WO or WO-like Islands (genomic islands that are associated with and/or derived from phage WO) demonstrated cytoplasmic incompatibility factors *cifA* and *cifB* as the primary cause of *Wolbachia*-induced CI and rescue [29–32]. In addition, transgenic expression of the WO-mediated killing gene *wmk* recapitulates male-specific embryo lethality and is a candidate for male killing [33]. Conversely, lytic activity of phage WO associates with reduced *Wolbachia* densities and CI levels [34].

First observed in 1978 as “virus-like bodies” within the gonads of *Culex pipiens* mosquitoes [35], phage WO is a temperate phage that exists in a lysogenic state (the integrated form of a phage genome is termed a prophage) until an event triggers particle production and subsequent lysis of the cell [4, 34, 36–38]. Unlike phages of free-living bacteria, however, the phage particles of intracellular *Wolbachia* contend with a two-fold cell challenge of bacterial and eukaryotic-derived membranes surrounding *Wolbachia* as well as the cytoplasmic and/or extracellular environments of the eukaryotic host. These unique challenges encountered by phage WO presumably selected for the evolution of a novel Eukaryotic Association Module (EAM) that comprises up to 60% of its genome with genes that are eukaryotic-like in function and/or origin [39]. The phage WO genome also features one of the longest genes ever identified in a phage and an abundance of ankyrin repeat domain genes [20, 23, 34, 40, 41], though their function has not been clearly elucidated as it has for the Ankyphages of sponge symbionts that aid in the evasion of the eukaryotic immune system [42]. Given the abundance and importance of phage WO in *Wolbachia* and for understanding genomic flux in endosymbioses worldwide, a firm grasp of its biology, including classification, evolution, and functions, will be important for establishing and comparing the rules across systems of endosymbiotic phages.

Here we survey prophage WO from 150 *Wolbachia* genome assemblies currently available in the NCBI database [43]. We report the patterns of distribution, chromosomal location, and functions of WO, and we propose a Linnaean classification system according to consultation with the International Committee and their guidelines on Taxonomy of Viruses (ICTV) [44, 45] in which there are three distinguishable phage WO genera within a new taxonomic order encompassing prophages of obligate, intracellular bacteria. We show that WO generally occurs in arthropod-associated *Wolbachia*, and prophage insertions are enriched away from the origin of replication in the bacterial chromosome. We fully annotate the EAM boundaries of representative WO genomes and highlight the presence of the CI genes, *cifA* and *cifB*, and a conserved set of eleven genes, defined here as the *Undecim Cluster*. We also establish a new model suggesting Octomom is derived from the EAM of prophage WO, with implications for Octomom-based pathogenicity, and we determine that all intact prophage WO genomes have a putatively novel patatin-based lytic cassette immediately upstream from the tail module. Finally, we report for the first time, to our knowledge, that prophage WO-like variants occur in diverse bacterial endosymbionts as well as metagenomes of putative symbionts from aquatic environments, providing a deeper understanding of WO origins, evolution, and ecology within and between endosymbiotic bacteria.

## Results

### Comprehensive survey of *Wolbachia’*s prophage WO and WO-like Islands

#### Prophage WO elements generally occur in arthropod-associated *Wolbachia*

*Wolbachia* occur in many protosome animal species of the superphylum Ecdysozoa, while prophage WO has previously been described as restricted to arthropod-associated strains. Because WO molecular surveys typically use single gene markers [15, 16], we comprehensively explored the NCBI database for prevalence of prophage WO, as determined by presence of one or more core phage WO genes (Fig 1a), throughout all sequenced *Wolbachia* genomes. All *Wolbachia* strains are indicated by a lower-case *w* followed by descriptor of host species, and prophage WO genomes are indicated by a WO prefix followed by the same host descriptor (listed in S1 Table).

**Fig 1.**
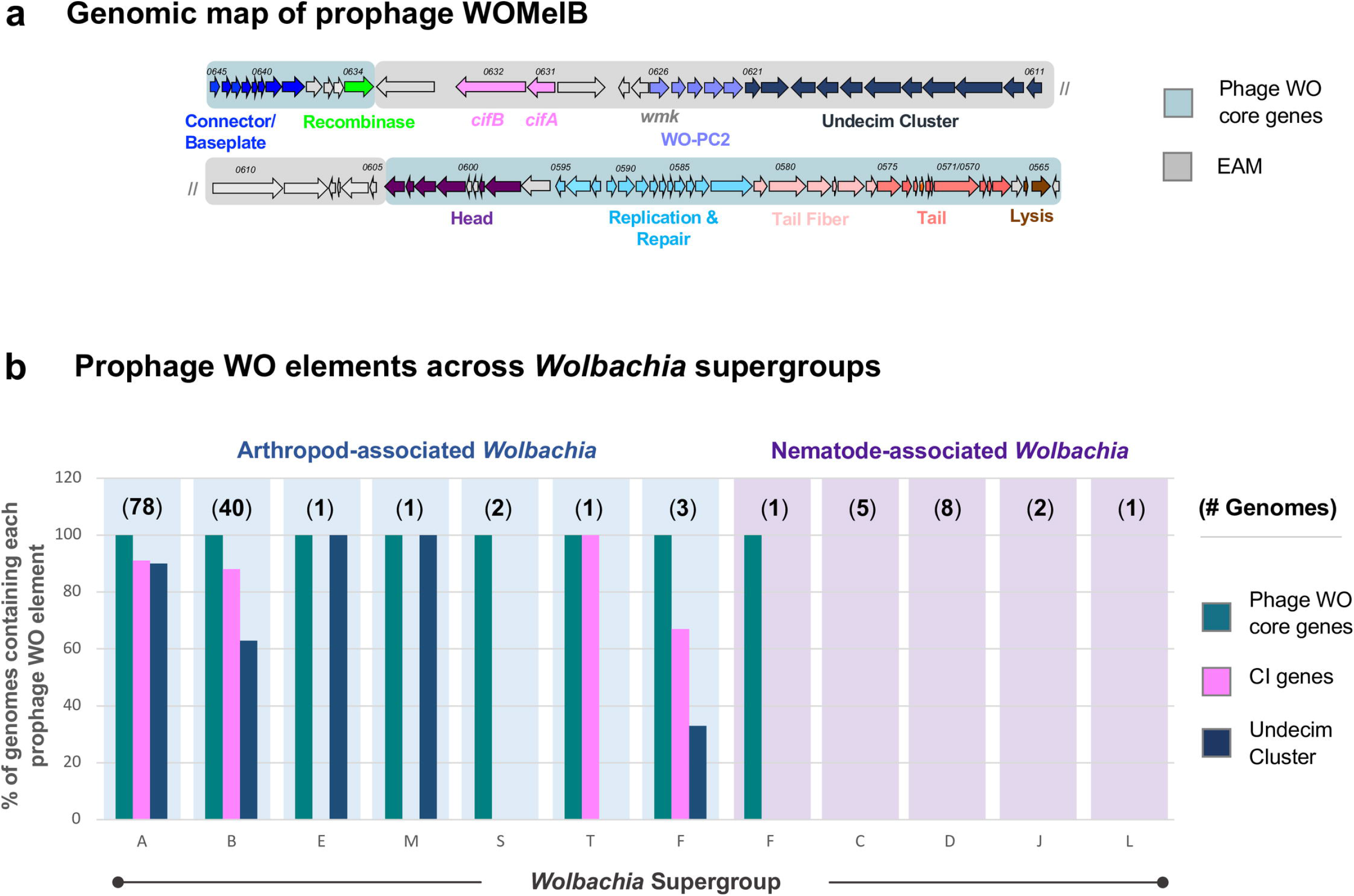
Prophage WO is modular in structure and associated with all arthropod-infecting *Wolbachia*. (a) A genomic map of prophage WOMelB from the *D. melanogaster w*Mel *Wolbachia* strain highlights phage WO core genes in blue and EAM genes in gray. Genes are illustrated as arrows and direction correlates with forward/reverse strand. The phage WO core consists of recombinase (green), connector/baseplate (royal blue), head (purple), replication and repair (light blue), tail fiber (light pink), tail (salmon), and lysis (brown). The WOMelB EAM encodes *cifA* and *cifB* (cotton candy pink), WO-PC2 containing HTH_XRE transcriptional regulators (lavender), and a conserved set of genes termed the *Undecim Cluster* (navy blue). (b) At least one phage WO core gene (teal) is associated with all sequenced arthropod-*Wolbachia* Supergroups and Supergroup F, which infects both arthropods (blue) and nematodes (purple). The Undecim Cluster (navy blue) is found in the majority of Supergroup A, B, E, and M *Wolbachia* genomes, and CI genes (pink) are encoded by the majority of Supergroup A, B, T, and F genomes. Phage WO elements are absent from all strictly-nematode *Wolbachia* Supergroups. The number of genomes analyzed is listed in parentheses above each Supergroup. Each bar indicates the % of genomes containing each phage WO element. Source data is provided in S1 Table.

Out of 150 assemblies across nematode and arthropod *Wolbachia*, phage WO occurs in arthropod *Wolbachia* with one exception from the mixed host supergroup of F *Wolbachia* (Fig 1b; S1 Table). All arthropod-associated strains contained evidence of intact or relic phage WO, termed WO-like Islands, and the single instance of WO genes in a nematode occurs in strain *w*Mhie from *Madathamugadia hiepei*, a parasite of the insectivorous South African gecko. The *w*Mhie genome encodes four genes that are conserved throughout phage WO’s transcriptional regulation and replication/repair modules (S2 Table) and are not part of the core *Wolbachia* genome. Interestingly, *w*Mhie is a member of Supergroup F that occurs in both arthropods and nematodes. Thus, the presence of phage WO genes in this *Wolbachia* genome supports a horizontal transfer of WO from arthropods to nematodes.

In addition to core phage WO genes, we characterized the widespread distribution of two phage WO elements across arthropod *Wolbachia*: (i) the cytoplasmic incompatibility factor genes *cifA* and *cifB* and (ii) Undecim Cluster (Fig 1b). Generally located within phage WO’s Eukaryotic Association Module (EAM [39]; Fig 1a) or in WO-like Islands (genomic islands that are associated with and/or derived from phage WO), *cifA* and *cifB* occur in Supergroups A, B, F, and T; the latter two are newly reported here. *Wolbachia* strains *w*Mov and *w*Oc of Supergroup F both encode phylogenetic Type I *cifA* and *cifB* genes, whereas *w*Chem of Supergroup T encodes Type II *cifA* and *cifB* genes (S3 Table; See [29, 46, 47] for a discussion of *cif* Types). Likewise, we identified a highly conserved set of eleven phage WO-associated genes, hereby termed the Undecim Cluster (Fig 1a, discussed below), that is distributed across most arthropod Supergroups but notably absent from all nematode *Wolbachia* genomes.

### Characterizing the prophage WO genome

#### Prophage WO genomes are comprised of conserved structural modules and a Eukaryotic Association Module

Prophage WO genomes adhere to the “modular theory” of phage evolution [18] and thus contain conserved structural gene modules (See discussion in S1 Text) and a Eukaryotic Association Module (EAM) [39]. To date, the EAM is unique to *Wolbachia*’s phage WO and as such is often overlooked by prophage prediction algorithms during the bacterial genome assembly process. Moreover, WO can markedly vary in gene content and synteny, and whether this variation does or does not sort into discrete genomic variants has not been investigated. Thus, we sought to identify conserved and distinguishing genomic features for a comprehensive nomenclature system for the community to classify phage WO major groupings. We mapped and re-annotated prophage WO regions from fully sequenced *Wolbachia* genomes to include the EAM and, more generally, incorporate updated annotations for each module.

All prophage WO regions were manually curated based on gene content and synteny (Fig 2; S1-S7 Figs) with regards to eight core phage modules (recombinase, replication & repair, head, connector/baseplate, putative tail fiber, tail, putative lysis, and EAM; labeled in Fig 1) and three newly identified and highly conserved gene clusters shown in Fig 2: (i) WO protein cluster 1 (WO-PC1), corresponding to hypothetical proteins WOCauB3_gp2-gp3; (ii) WO protein cluster 2 (WO-PC2), located within the EAM and corresponding to putative HTH_XRE transcriptional regulators, DUF2466 (formerly RadC), and hypothetical proteins WOMelB_WD0622-WD0626; and (iii) the Undecim Cluster, an eleven-gene region located within the EAM and corresponding to WOMelB_WD0611-WD0621.

**Fig 2.**
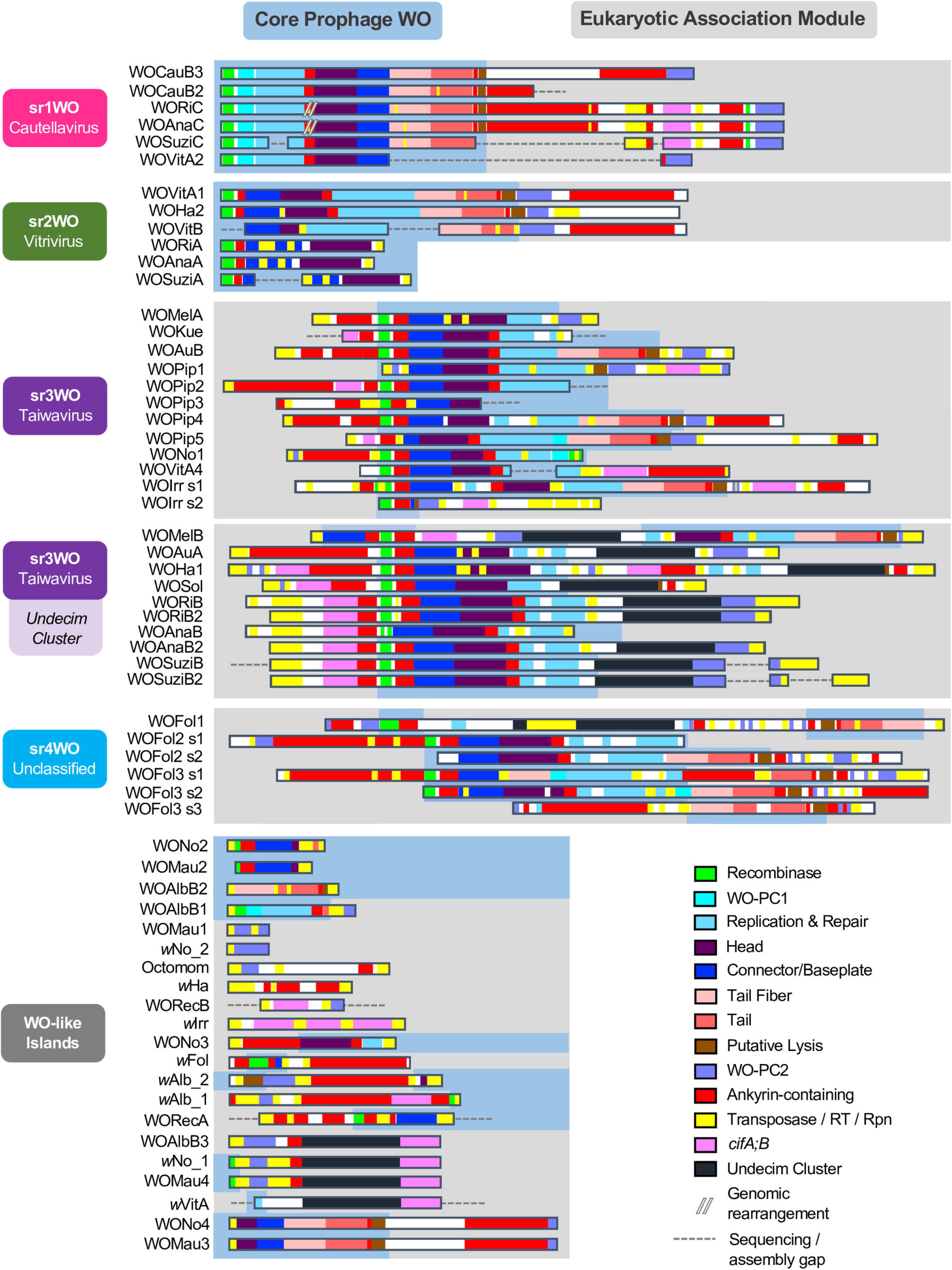
Prophage WO variants feature distinguishable module synteny. Prophage WO variants are organized by genome content and synteny of their structural modules. Sr1WO and sr2WO feature a 5’-core prophage WO region (blue) and a 3’-EAM (gray). Sr3WO features an internal core prophage WO region that is flanked by EAM genes and mobile elements (yellow). Sr4WO is only present in *w*Fol and features three genomic regions with multiple prophage segments. WO-like Islands feature small clusters of prophage WO-like genes; they are comprised of singular structural modules and/or subsets of EAM genes. All modules are color coded: green = recombinase; turquoise = WO-PC1; light blue = replication; purple = head; blue = connector/baseplate; light pink = tail fiber; salmon = tail; brown = putative lysis; lavender = WO-PC2; and navy blue = Undecim Cluster. In addition, ankyrins are shown in red; transposable elements are shown in yellow; and *cifA;cifB* are shown in cotton candy pink. Dotted lines represent breaks in the assembly; module organization is estimated based on closely related variants. Sr1WO is highlighted in hot pink; sr2WO is highlighted in green; sr3WO is highlighted in purple; sr4WO is highlighted in blue; WO-like Islands are highlighted in gray.

#### There are four distinguishable prophage WO variants: sr1WO, sr2WO, sr3WO, and sr4WO

While gene synteny within each core module is generally consistent, the arrangement of modules across prophage genomes is variable and does not correlate with the early organization of *orf7*-based WO clades, WO-A and WO-B [16, 48]. To formally update this classification with a more comprehensive classification system, we identified conserved WO loci and modular synteny diagnostic of the four WO arrangement groupings that reflect genus-level ranking. Sequence variation in one gene candidate was consistently associated with similar variation in gene content and synteny: the large serine recombinase [18, 49]. Phage-encoded large serine recombinases facilitate integration of the phage genome into specific attachment sites within the bacterial chromosome as well as control the excision, often with the help of an accessory protein, of the prophage genome during the lytic cycle [50]. A BLASTN analysis of the WO serine recombinase gene confirmed that only those associated with comparable WO module arrangement were full-length reciprocal BLAST hits. Phylogenetic analysis of the recombinase peptide sequence also supported four distinct genus-level clades of prophage WO (common names sr1WO, sr2WO, sr3WO, and sr4WO; nomenclature proposed in [49] and based on the “<u>s</u>erine recombinase”) as well as closely-related recombinases in prophage regions of non-*Wolbachia* endosymbionts, including the *Paramecium* endosymbiont *Holospora obtusa* (Fig 3a). The genomic content, organization, and chromosomal integration of each srWO variant are described below.

**Fig 3.**
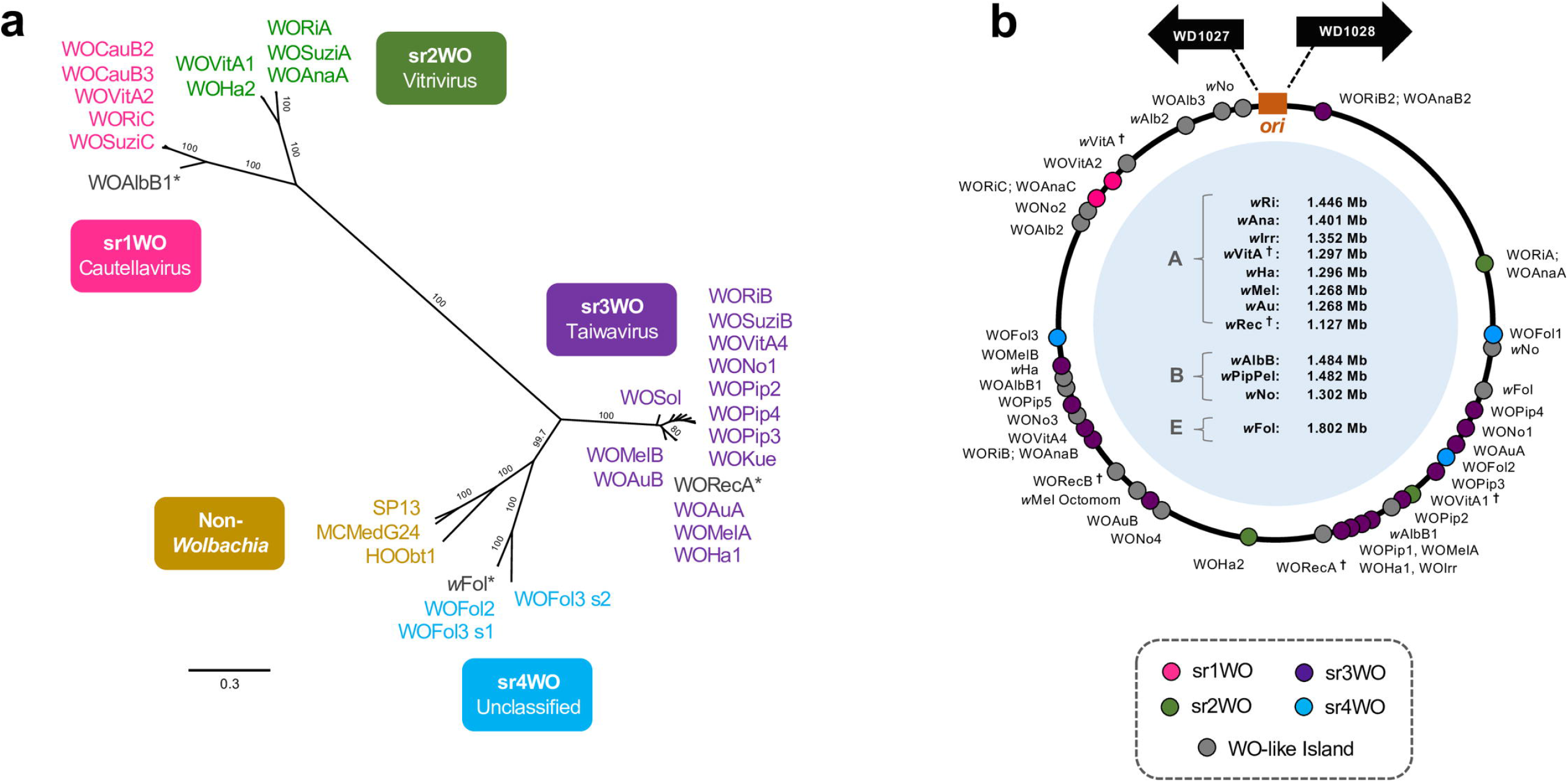
Phylogeny of prophage WO’s large serine recombinase correlates with module synteny and genomic integration. (a) A phylogenetic tree of prophage WO’s recombinase sequence illustrates the utility of this gene as a WO-typing tool to classify prophage WO variants. Four distinct clades correlate with sr1WO-sr4WO genome organization shown in Fig 2. Non-*Wolbachia* sequences represent similar prophages from other bacterial hosts, such as the prophage HOObt1 of *Holospora obtusa*, an endonuclear symbiont of *Paramecium*. The tree was generated by Bayesian analysis of 283 amino acids using the JTT-IG model of evolution. Consensus support values are indicated for each branch. (*) indicates that the prophage regions are highly degraded; while they likely originated from the corresponding prophage group, they are now classified as WO-like Islands (S7 Fig). (b) Prophage WO integration loci are concentrated opposite the origin of replication, *ori*. All *Wolbachia* genomes have been standardized where each dot represents % nucleotide distance calculated by: (nucleotide distance between 5’-WO and *ori* / genome size) * 100. (**^†^**) indicates the genome is not closed/circularized; genomic locations are estimated based on alignment of contigs to a reference genome (obtained from authors in [51, 52]).

*sr1WO*. The proposed genus-level taxonomic name for sr1WO, described below, is Cautellavirus. Most sr1WO recombinases integrate into *Wolbachia*’s magnesium chelatase gene, as we previously reported [39], with portions of the bacterial gene found flanking either side of the prophage region. Two exceptions are in: (i) closely-related *w*Ri and wAna where the sr1WO prophage has since been rearranged in the *Wolbachia* genome (S1 Fig) with a portion of the magnesium chelatase now associated with each prophage fragment (S8a-b Fig); and (ii) *w*CauB which contains at least two sr1WO prophages, and WOCauB3 has a secondary intergenic attachment site between *sua5* and a hypothetical protein (S8c Fig).

A key characteristic of sr1WOs is the single domain HTH_XRE transcriptional regulators of WO-PC2 (S1 Fig, lavender) that are located at the 3’-end of the prophage region. Because the genes are fused in most other WO prophages, they are sometimes annotated as pseudogenes (i.e., wRi_p006660 and wRi_p006630 of WORiC) in the *Wolbachia* genome; however, conservation across multiple variants suggests they are functional. Sr1WOs also lack the methylase/ParB gene that is associated with all other WO prophages. A few genomes (i.e, WORiC, WOAnaC, WOSuziC) harbor *cifA* and *cifB* genes, though the origin of these genes remains inconclusive due to a downstream, highly-pseudogenized sr3WO recombinase (wRi_p006680) and adjacent transposases. Finally, all members of the sr1WO group have a distinct 5’-core-prophage region followed by an ankyrin-rich 3’-EAM (Fig 2 and S1 Fig).

*sr2WO*. The proposed genus-level taxonomic name for sr2WO, described below, is Vitrivirus. sr2WO prophages genes are also organized as 5’-core-prophage followed by 3’-EAM (Fig 2 and S2 Fig), yet module synteny is quite distinct from sr1WO: (i) they lack WO-PC1; (ii) the replication, head, and connector/baseplate modules are reversed; (iii) WO-PC2 is located at the juncture between the core-prophage and EAM regions rather than at the terminal 3’-end of the prophage genome; and (iv) *cifA* and *cifB* genes are absent from assembled genomes thus far. The sr2WO recombinase integrates into variable number tandem repeat 105 (VNTR-105) as previously reported [39], a conserved intergenic region used to type closely-related A-*Wolbachia* strains [53]. While flanking, disrupted portions of the magnesium chelatase correlate with prophage boundaries of sr1WO genomes, disrupted VNTR-105 regions likewise flank the complete sr2WO genome, including the eukaryotic-like *secA* [54] EAM of WOHa2.

*Sr3WO*. The proposed genus-level taxonomic name for sr3WO, described below, is Taiwavirus. Unlike the previous groups, sr3WO appears to lack a conserved integration site. Rather, these variants feature a core prophage region that is flanked on either side by EAM regions, are separated from adjacent *Wolbachia* genes by an enrichment of transposase-encoding insertion sequences (Fig 2, yellow and S4 Table), and are concentrated away from the origin of replication in the bacterial chromosome (Fig 3b). While their function here is unknown, transposable Mu-like phages replicate via replicative transposition in the bacterial chromosome and, much like phage WO, are associated with severe chromosomal rearrangements and disruptions [55]. Under a similar model, sr3WO transposases could mediate prophage replication and movement throughout the *Wolbachia* genome.

Sr3WO core-prophage module synteny generally resembles that of sr2WO, although a subset of variants also encode an eleven-gene module termed the *Undecim Cluster* (S4 Fig and S5 Fig), discussed in detail below. Most importantly, unlike other prophage WO groups, a majority of the sr3WO variants contain at least one *cifA* and *cifB* gene pair, the locus responsible for *Wolbachia*’s cytoplasmic incompatibility phenotype [29, 30, 32, 46, 47].

*Sr4WO*. The prophage WO group identified strictly in *w*Fol of *Folsomia candida* springtails is tentatively labelled sr4WO. Unlike the above clades, sr4WO will remain unclassified at the genus level due to high variability and rearrangement of the prophage genomes. A formal classification will be evaluated when more genomes are sequenced that support conserved taxonomic characteristics for the clade. Three variants, broken into multiple segments (S6 Fig), loosely resemble the module synteny of sr3WO. WOFol1 is associated with an Undecim Cluster similar to sr3WO, but all variants contain single-domain HTH_XRE genes similar to sr1WO. The sr4WO prophages contain multiple genomic duplications and mobile elements [56]. While they appear to lack *cifA* and *cifB* genes, they are enriched with multiple copies of *ligA* and resolvase. More variants of this group are needed to analyze chromosomal integration.

#### WO-like Islands

We identified numerous portions of the prophage WO genome that do not contain enough genetic information to be properly classified. Termed WO-like Islands, they are comprised of single core phage modules, such as a baseplate or tail, and/or genes that are typically associated with the prophage WO genome rather than part of the core *Wolbachia* genome (Fig 2 and S7 Fig). Most WO-like Islands are therefore considered “cryptic”, “relic”, or “defective” prophages, and likely originated from an ancestral prophage WO genome where they have since been domesticated by the bacterial host or are in the process of degradation and elimination from the chromosome. Based on studies in other systems, conserved prophage genes or gene modules that are not part of a complete prophage are likely to provide a fitness advantage to their host [57, 58] and may interact with, even parasitize, fully intact phages within the same bacterial host [59, 60].

Like sr3WO prophages, WO-like Islands are often flanked by at least one insertion sequence (S4 Table) and are commonly associated with CI genes *cifA* and *cifB*. In the unusual case of the *w*Irr WO-like Island, four CI loci, along with multiple transposases, are arranged in a single genomic cluster that is not associated with conserved WO genes (S7 Fig). We tentatively label the region as a WO-like Island because (i) the *cif* genes and adjacent hypothetical proteins are overwhelmingly associated with prophage WO regions and (ii) there is evidence of a highly disrupted prophage genome about 160kb upstream in the *w*Irr chromosome (S4 Fig) that is also enriched with transposases, allowing for the possibility of a prophage WO origin. Such a model for the putative phage WO origin of one highly studied WO-like Island, *w*Mel’s Octomom, is discussed in detail below.

#### Prophage WO is spatially concentrated away from the origin of replication in the *Wolbachia* chromosome

To comprehensively examine the association of each prophage WO variant with its chromosomal location in *Wolbachia*, we mapped integration sites, determined by the recombinase or the most 5’-WO gene, on the chromosome with respect to normalized distance from the putative origin of replication, *ori* [61]. There is a clustering of prophage WO insertion loci, particularly sr3WOs, opposite the origin of replication (Fig 3b; Chi-square 2-tailed, p=0.0035) that is similar to the localization patterns of temperate phages in *Escherichia, Salmonella*, and Negativicutes [62–65]. WO chromosomal location patterns support a model in which prophage insertions and WO-like Islands may not be tolerated in regions directly surrounding the origin of replication.

#### Transposable elements may facilitate transposition and domestication of prophage WO regions

In addition to specific chromosomal integration patterns, we next surveyed the relationship between WO and its associated mobile elements. With the exception of WOCauB3, all fully sequenced prophage WO genomes and WO-like Islands contained at least one transposable element beyond the phage recombinase. The diversity of the WO-associated transposable elements by prophage variant is listed in S4 Table and includes (i) transposases of insertion sequence families IS3, IS4, IS5, IS6, IS66, IS110, IS256, IS481, IS630, IS982; (ii) recombination-promotion nuclease (Rpn), which encodes a PD-(D/E)XK nuclease family transposase; and (iii) reverse transcriptase of group II intron origin (RT). WO’s transposable elements are associated with the genomic rearrangement (e.g., WORiC), degradation or domestication (e.g., WORiA), and copy number variation (e.g., WORiB) of various prophage genomes. As discussed above, flanking transposases of sr3WO variants may also play a role in replicative transposition similar to phage Mu.

We observed that reverse transcriptases of group II intron origin (RT) are associated with chromosomal rearrangements, insertions, and/or duplications of multiple sr3WO and sr4WO prophages (illustrated in S9 Fig). Likewise, we identified numerous associations of *cifA;B* gene pairs with RTs of sr3WO variants (including WOPip1, WOVitA4, WOIrr, WOHa1, WORiB, WOAnaB, WOSuziB) and the *w*Irr WO-like Island. Therefore, the association of CI loci with transposable elements – both within and beyond prophage regions – could be indicative of post-integration genomic rearrangement and/or domestication of the genes, as previously discussed [6]. Below we propose a detailed model and evidence for the most intriguing RT-associated genomic rearrangement, the origin of *w*Mel’s Octomom from prophage WOMelA to generate a WO-like Island (Fig 4).

**Fig 4.**
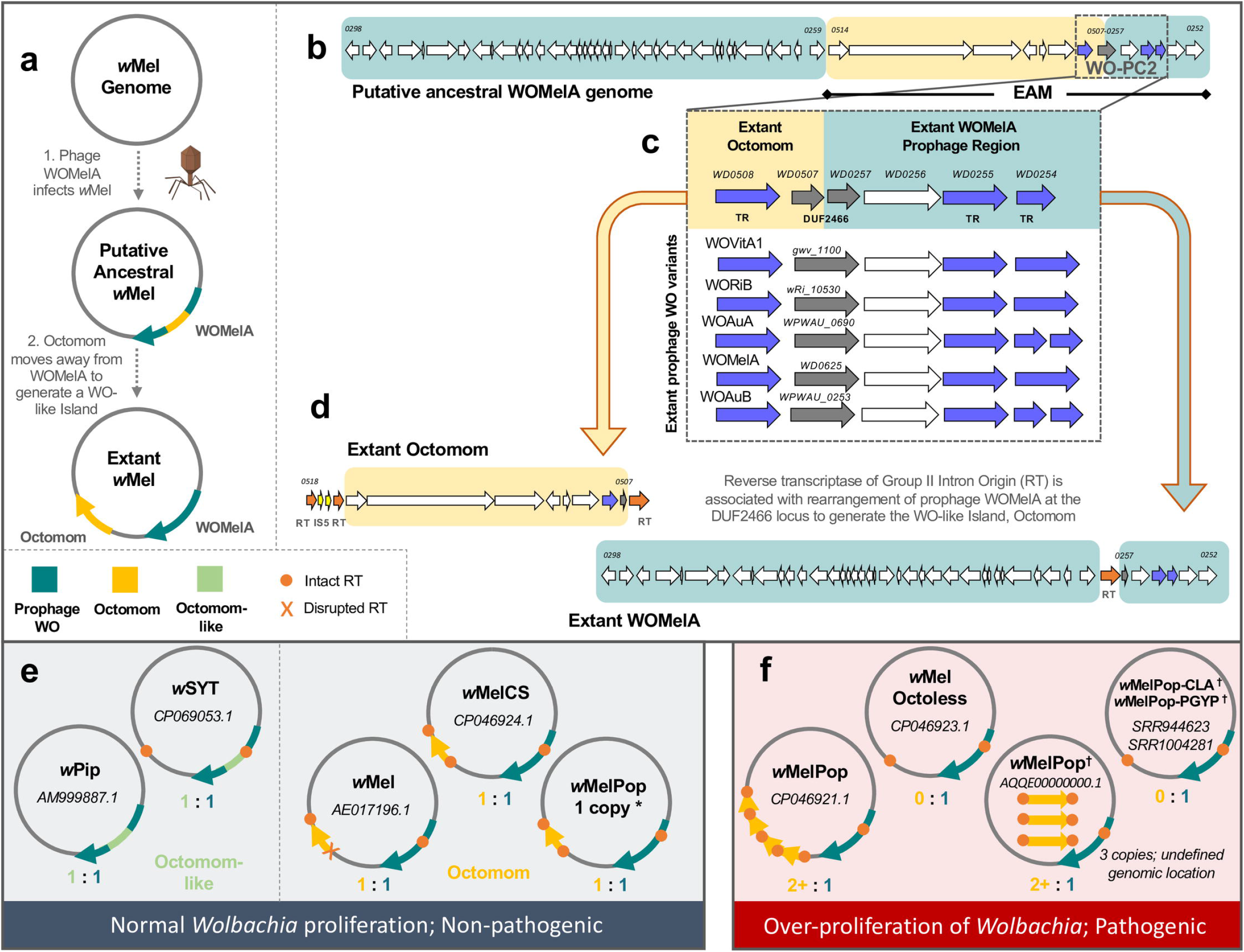
Comparative genomics supports a WO:Octomom origin model for *Wolbachia* proliferation in wMelPop. (a) A new model for Octomom origin predicts the initial infection of *w*Mel with a WOMelA phage. After integration, Octomom splits from the WOMelA core prophage region to form a WO-like Island. (b) A genome map of the putative, intact, ancestral WOMelA where Octomom is highlighted in yellow and the extant WOMelA genome in teal illustrates placement of Octomom in the WO EAM. (c-d) An alignment of the WO-PC2 region with closely related prophages shows that half of the conserved module (WD0507-WD0508) is now associated with Octomom and the other half (WD0257-WD0254) remained with WOMelA prophage region. DUF2466 is split across the genomic regions and, when concatenated, shares homology to intact DUF2466 genes of WO-PC2. An IS5 insertion (d) is associated with single-copy Octomom stability in the *w*Mel chromosome. In wMelCS-like genomes, where the flanking RTs are intact (see S10 Fig), Octomom varies in copy number. (e) When Octomom (orange-yellow) and Octomom-like (green, defined by homology to WD0512, WD0513 and WO-PC2 and illustrated in S10 Fig) regions exist in a single copy, either within or outside the corresponding prophage region, *Wolbachia* proliferation is normal, and it is non-pathogenic. (f) If the WO-like Island occurs in multiple copies or is absent from the genome, *Wolbachia* over-proliferate and are pathogenic. (*) Restoring the 1:1 (WO:Octomom) ratio returns the *w*MelPop phenotype back to normal levels. The association of Octomom with pathogenicity (i.e., correlation vs. causation) is still to be determined [66–68]. NCBI accession numbers are listed for each genome; (†) indicates circular genomes are unavailable and genomic locations are putative.

### Unique characteristics of prophage WO

#### The WO-Octomom Model posits that Octomom is derived from the EAM; *Wolbachia* proliferation may be dependent upon a 1:1 ratio of Octomom : prophage WO

Octomom is a cluster of eight genes in the *D. melanogaster w*Mel *Wolbachia* genome that has been described for its resemblance to a bacterial pathogenicity island (see S10 Fig for genome schematic) [69]. Increasing the environmental temperature of flies either containing multiple copies or completely lacking this region results in *Wolbachia* over-proliferation and pathogenicity [67, 68]. Based on our observations of RT-associated genomic rearrangement, we present a new WO-Octomom Model (Fig 4a) with genomic evidence (Fig 4b-d), in which Octomom putatively originated from the EAM of ancestral WOMelA (sr3WO). First, an ancestral phage WOMelA with core phage genes as well as an Octomom-encoding EAM infects *w*Mel and integrates into the bacterial chromosome. Second, Octomom splits from the prophage EAM region, possibly mediated by RTs, to form an independent WO-like Island about 38kb from the extant WOMelA (Fig 4a). This is supported by gene synteny of the WO-PC2 variant that is split between Octomom and WOMelA at the DUF2466 gene (also annotated as *radC*). Notably, by concatenating the two regions at Octomom’s WD0507 (5’-DUF2466) and WOMelA‘s WD0257 (3’-DUF2466), the gene synteny forms a complete WO-PC2 and closely resembles that of related sr3WO prophages (Fig 4b-d).

Furthermore, Octomom homologs of the two-domain HTH_XRE transcriptional regulator (WD0508) are characteristic of sr2WO and sr3WO prophages, and the *mutL* paralog (WD0509) from Octomom is a phage WO-specific allele [70] that is distinct from the chromosomal *mutL* (WD1306). This supports an ancestral WOMelA prophage genome comprised of core structural modules and an Octomom-containing EAM with intact WO-PC2 (Fig 4b). An alternative explanation could be that genes WD0512-WD0514 existed as a pathogenicity island in the *Wolbachia* chromosome prior to WOMelA infection and later acquired adjacent EAM genes from the prophage to form a complete Octomom Island. In this case, we would expect to find at least one other instance of WD0512-WD0514 occurring independent of prophage regions in other *Wolbachia* strains. Instead, the only *Wolbachia* homologs, to date, are associated with the EAMs of WOPip5 and the wSYT (*Wolbachia* of *Drosophila santomea, D. yakuba*, and *D. teissieri*, respectively) prophages [6, 19, 71] (S10 Fig).

An interesting and robust correlation of this WO-Octomom Model is that one copy relative to prophage WO, either within or outside of the prophage region, is always a distinguishing factor of non-pathogenic *Wolbachia* (Fig 4e), while absence *or* multiplication of Octomom are notably associated with *Wolbachia* over-proliferation and pathogenicity (Fig 4f). This has been previously reported in context of the *Wolbachia* chromosome [66, 67], and we make the distinction here of a *prophage* association to enable a more fine-tuned exploration of Octomom biology. For example, the disruption (*w*Mel) or absence of one (*w*SYT) or both (*w*Pip) flanking RTs correlates with a static 1:1 ratio of the Octomom-like region (i.e., containing WD0512-WD0513 and a transcriptional regulation gene) and its corresponding prophage genome (Fig 4e). Conversely, the region is flanked by identical RTs on either side in all *w*Mel clade VI strains, including *w*MelCS and the dynamic *w*MelPop that ranges from 0 to multiple copies of the WO-like Island (Fig 4f; *w*Mel phylogeny presented in [66, 72]). When the 1:1 ratio in clade VI strains is disrupted, possibly in conjunction with flanking RTs, *Wolbachia* develops a pathogenic relationship with its animal host [66, 72]. The possible association of RTs with Octomom copy number is also notable due to the observed dependence of both RT activity [73, 74] and *w*MelPop pathology [67, 68] on environmental conditions, such as temperature. The direct role of Octomom on host phenotype is a subject of debate [66, 67], and understanding the association of prophage WO with this region, if any, could inform the biology of this unique system. The two phage-derived regions, for example, may share a common regulatory mechanism since the proposed ancestral splitting of Octomom from WOMelA broke a cluster of transcriptional regulators, namely one transcriptional regulator (WD0508) from the other two (WD0254 and WD0255) that would typically form an intact module. Alternatively, a split of Octomom from its associated prophage genome may influence epigenetic modifications via WOMelA’s adenine methylase (WD0267; see [66] for a discussion of epigenetic vs. genetic factors).

#### Undecim Cluster is a unique eleven gene island associated with prophage WO

Another “pathogenicity island” candidate in the *Wolbachia* chromosome is a highly conserved set of genes (WD0611 to WD0621; Fig 5a) defined here as the *Undecim Cluster (Undecim* is Latin for “eleven”). We identify it in the majority of WO-containing *Wolbachia* genomes (Fig 1b), particularly in association with *cifA-*and *cifB*-encoding regions of sr3WO (S4 Fig and S5 Fig) and WO-like Islands (S7 Fig). Unlike sr3WO prophages themselves, however, the Undecim Cluster does not occur more than once per *Wolbachia* genome. Its complete absence from both *w*Pip and *w*Rec suggests that it is not strictly required for *Wolbachia*’s intracellular survival and/or ability to induce cytoplasmic incompatibility. Rather, it may contribute to variation in host-symbiont interactions [18, 48] by encoding a broad spectrum of metabolic functions and transport potential [75, 76], including cellular exopolysaccharide and/or lipopolysaccharide (LPS) biosynthesis (WD0611-WD0613; WD0620), methylation (WD0613-WD0614; WD0621), production and export of antibiotics and cytotoxic compounds (WD0615-WD0616) and metabolite transport and biosynthesis (WD0617-WD0619) (Fig 5b). It was identified in phage particle genomes from both *w*VitA and *w*CauB [39], indicating that the region may be transferred between *Wolbachia* strains via the phage. In addition, both RNA-SEQ [77] and mass spectrometry data [75] show that the region is highly expressed. Interestingly, ten of the eleven genes were involved in a lateral gene transfer event between *Wolbachia* and the *Rickettsia* endosymbiont of *Ixodes scapularis* (REIS; [17, 76]) with WD0612 to WD0618 sharing 74% nucleotide identity to a region of the Rickettsial plasmid pREIS2 and WD0619 to WD0621 sharing 67% identity to a region of the bacterial chromosome (Fig 5a). We also identified homologs in *Cardinium hertigii* cHgTN10 (CP029619.1; 67% nucleotide identity) and *Phycorickettsia trachydisci* (CP027845.1; 68% nucleotide identity). While not contiguous in *C. hertigii*, adjacent transposases may have facilitated post-integration rearrangement.

**Fig 5.**
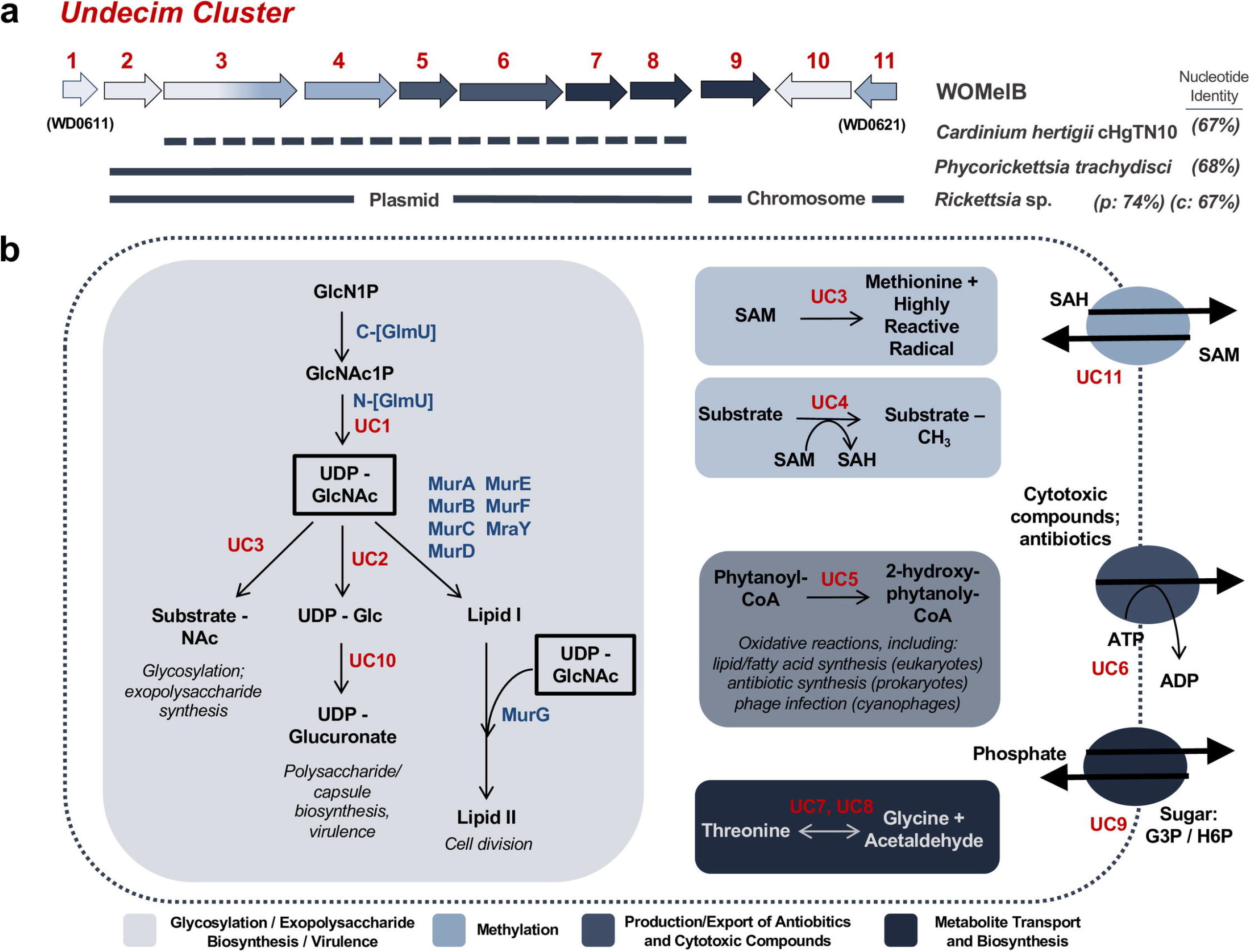
The Undecim Cluster contributes a wide range of cellular processes associated with host-symbiont interactions. (a) A genome map illustrates prophage WO’s Undecim Cluster. Gene labels UC1 -UC11 correlate with *w*Mel locus tags WD0611-WD0621. Lines under the genes indicate lateral gene transfer events of this region between *Cardinium hertigii* cHgTN10, *Phycorickettisa trachydisci*, and multiple strains of *Rickettsia*, including the *Rickettsia* endosymbiont of *Ixodes* scapularis (REIS) and its plasmid (pREIS2). Nucleotide identity is listed to the right. Dashed lines indicate that the region is not contiguous in the genome. UC1 shares partial homology with a core *Wolbachia* gene, glmU (WD0133) and was either not involved in the transfer event or has since been lost from non-*Wolbachia* genomes. (b) A cellular model illustrates the putative functions associated with this region. Cellular reactions are highlighted in boxes and membrane transporters are drawn as ovals. *Wolbachia* genes are labeled in blue; Undecim Cluster genes are labeled in red. UC3 (WD0613) is a fusion protein with an N-terminal glycosyltransferase and C-terminal radical SAM domain; therefore, it is listed twice. Reactions in light gray are likely precursors to multiple pathways in glycosylation, exopolysaccharide biosynthesis, cell division, and/or virulence. Light blue is associated with methylation; dark gray is associated with the production and export of antibiotics and cytotoxic compounds; and navy blue is associated with metabolite transport and biosynthesis. The above functions are predicted based on annotation and homology to other systems. Given the contiguous conservation of the Undecim Cluster throughout prophage WO, all functions, including those not captured in this model, are likely interrelated and influence host-symbiont dynamics.

#### Phage WO putatively harbors a novel lytic cassette

The most direct impact on *Wolbachia* cellular biology is the potential for phage WO to induce cell lysis [34, 78]. The mechanism of phage-induced cell lysis has been well documented and generally involves a three-component lysis system in gram-negative infecting phages: endolysin, holin, and spanins [79]. This genetic system is noticeably absent from prophage WO genomes, and peptidoglycan, the bacterial target of canonical phage endolysins, has never been detected in *Wolbachia* [80]. We therefore hypothesized that WO phages encode an alternative lytic pathway. The top candidate is a putative and novel patatin-based lytic cassette immediately upstream from the tail module [81].

The cassette contains a patatin-like phospholipase A_2_, a small holin-like protein, and an ankyrin-repeat protein. A few prophage WO variants (i.e., WOVitA1, WOAuB, WOPip1, WOPip4, and WOPip5) additionally encode an endonuclease of the phospholipase D family. Patatin-like proteins determine virulence in multiple gram-negative bacteria and specifically facilitate disruption of host cell membranes by *Pseudomonas aeruginosa* and *Rickettsia typhi* [82, 83]. They are significantly more common in pathogenic bacteria and symbionts than in non-pathogens, suggesting a role in host-association [84]. Holins are not easily annotated because they do not share conserved domain sequence homology, yet several lines of evidence suggest the small protein adjacent to patatin is a “holin-like” candidate: it (i) encodes a single N-terminal transmembrane domain with no predicted charge; (ii) features a C-terminal coiled coil motif; (iii) is smaller than 150 amino acid residues; and (iv) has a highly charged C-terminal domain (S11a Fig) [79, 85, 86]. In addition, homologs of this holin-like gene in prophages from bacterial chromosomes other than *Wolbachia* (e.g., a Tara Oceans Prophage and *Holospora* sp.) are directly adjacent to a GH108 lysozyme, further supporting its holin-like potential (S11b and S11c Fig, Fig 6). The third conserved gene in this module, an ankyrin repeat protein with a C-terminal transmembrane domain, may have the potential to impact membrane stability similar to spanins of the traditional phage lysis model; alternatively, they may play a role in evasion of the arthropod-host immune response similar to those in sponge-associated Ankyphages [42]. Together, this module is fairly conserved across tailed WO phages and is a likely candidate in the exit and/or entry of phage particles through *Wolbachia*’s multiple membranes.

**Fig 6.**
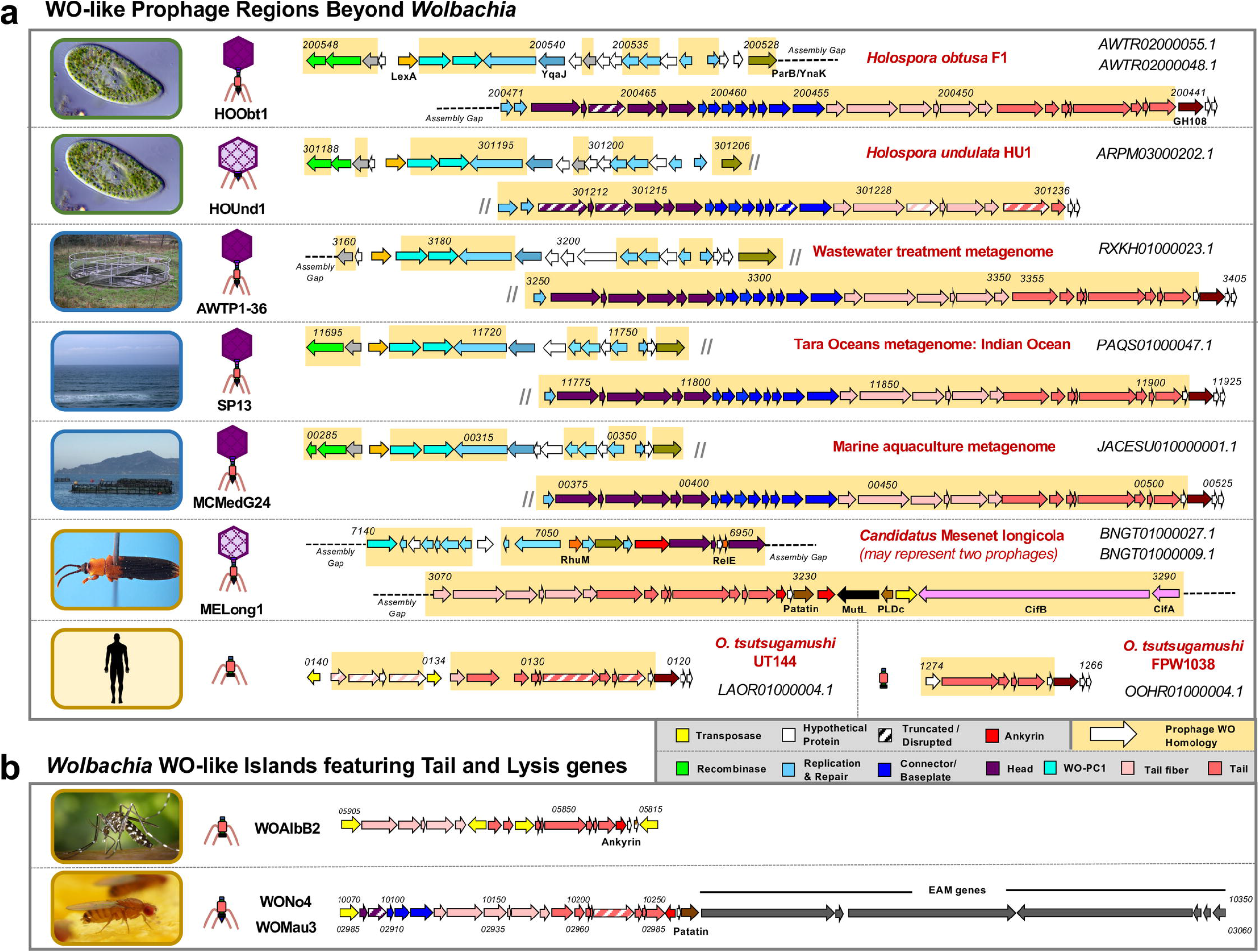
WO-like prophage regions are found in endonuclear *Paramecium* endosymbionts, aquatic environments, and other animal-associated bacteria. (a) Genome maps of non-*Wolbachia* prophage regions illustrate similar gene content and synteny to prophage WO. Locus tags are listed in italics above the genes; NCBI contig accession numbers are shown in the right-hand corner of each genome. Dashed lines represent breaks in the assembly whereas small diagonal lines represent a continuation of the genome onto the next line. Genes with nucleotide homology to prophage WO are highlighted in yellow and genes of similar function are similarly color-coded according to the figure legend. *Candidatus* Mesenet longicola is the only genome to feature EAM genes, including *cifA* and *cifB*. Arrows with diagonal stripes represent genes that may be pseudogenized relative to homologs in other prophage genomes. Genome maps for *H. elegans and H. curviuscula* prophages are not shown. (b) WO-like Islands featuring tail and lysis genes share homology with the *Orientia* regions and may represent phage-derived bacteriocins. Predicted physical structures are illustrated to the left of each genome. Images illustrate the isolation source for each prophage: green borders represent protozoa; blue borders represent aquatic environments; gold borders represent animals.

#### Other prophage genes in the *Wolbachia* chromosome are Gene Transfer Agents (GTAs)

In addition to prophage WO, we identified several non-WO prophage genes (S12 Fig) in the majority of *Wolbachia* Supergroups, including those of the filarial nematodes. Similar to the well-studied GTA of *Rhodobacter capsulatus* (RcGTA; [87, 88]), at least six of these genes encode *E. coli* phage HK97-like conserved domains (S5 Table). We also identified GTA terminase genes associated with the *Wolbachia* chromosome. As reported for *Rickettsiales*, the GTA loci are found in multiple locations across the genome rather than organized in an identifiable prophage-like cluster [89]. To investigate the evolutionary relationship of the GTA genes with their *Wolbachia* host, we performed individual nucleotide alignments and recovered two highly conserved genetic groups that demarcate Supergroup A and B *Wolbachia* (S13 Fig), supporting vertical descent with modification across these major supergroups. While absent from Supergroups J and L of nematodes, they are present across all other *Wolbachia* Supergroups as well as the closely related genera *Candidatus* Mesenet*, Anaplasma, Ehrlichia*, and *Rickettsia* (S12b Fig). These results imply that *Wolbachia*’s GTA genes are vertically inherited, codiverge with their bacterial hosts, and likely functional given their intact sequences. They are, however, distinct from phage WO, not indicative of former WO-infections, and may be lost during genome reduction.

### Prophage WO beyond *Wolbachia*

#### Prophage WO-like variants occur in diverse bacterial endosymbionts and metagenomes

We identified multiple prophage WO-like variants beyond the *Wolbachia* genus that have gene synteny and nucleotide identity to prophage WO structural modules in: (i) endonuclear bacterial symbionts of *Paramecium* (*Holospora obtusa, H. undulata, H. elegans*, and *H. curviuscula*) [90]; (ii) metagenome projects from an advanced water treatment facility [91], the Indian Ocean (*Tara* Oceans circumnavigation expedition [92]), and a marine aquaculture habitat [93]; (iii) *Candidatus* Mesenet longicola, the CI-inducing bacterial endosymbiont of *Brontispa longissima* [94]; and (iv) multiple strains of *Orientia tsutsugamushi* isolated from humans (Fig 6a). While the structural genes closely resembled those of prophage WO, novel genes were identified in the replication/repair and lysis modules (Fig 6a, genes with prophage WO homology are highlighted in yellow). All non-*Wolbachia* variants except *Candidatus* Mesenet longicola lacked signature *Wolbachia* phage WO genes such as patatin, ankyrin repeats, and the EAM that are putatively or definitively involved in phage-by-arthropod interactions.

Relative to the full-length genomes recovered from *Holospora*, *Candidatus* Mesenet longicola and the metagenome projects, *Orientia* prophages appeared to be highly degenerate. These regions featured only tail and lysis genes, but the modules are noticeably intact. Some WO-like Islands, such as WOAlbB2, WONo4, and WOMau3 (Fig 6b), also harbor sole tail and lysis modules. The retention of a complete phage structural module in the bacterial chromosome suggests that it has been domesticated and adapted to benefit the host. For example, several studies report phage-derived bacteriocins that consist of tail and lysis genes and target other strains of the same bacterial species [57]. Similarly, an extracellular contractile injection system (eCIS) comprised of phage tail-like proteins specifically targets eukaryotic cells [95]. Overall, the presence of WO-like variants in non-*Wolbachia* genera continue to support phage WO lateral transfer between unrelated, coinfecting symbionts. This is further evident by the presence of the CI genes, *cifA* and *cifB*, in the *O. tsutsugamushi* genome [96], which may represent a derived variant of phage WO from *Wolbachia* that has since been domesticated by its bacterial host. Alternatively, the association of CI genes in a bacterium harboring WO-like variants could be indicative of two other possible origins - either the last common ancestor of the WO and WO-like phages encoded *cifA* and *cifB*, or the loci may have originated in WO-like phages and transferred to *Wolbachia*. For divergent, horizontally transferred elements, it is often not possible in practice to assign a direction of evolution and origin story.

### Linnaean classification of phage WO

Finally, while phage WO is a model organism to study the tripartite association between viruses, endosymbiotic bacteria, and animal hosts, it is not yet recognized by the International Committee on Taxonomy of Viruses (ICTV). Recently, the ICTV Executive Committee implemented a pipeline for the official classification of viruses from metagenomic datasets [45], including those originating from integrated prophage sequences. Through our comparative analysis of prophage WO sequences here with those that have been sequenced from active particles (i.e., WOVitA1 and WOCauB3), we propose a formal phage WO taxonomy (Fig 7) to align with the ICTV Linnaean-based classification code [44]. The correlation between common name and proposed scientific name for each taxonomic rank is listed in Table 1.

**Fig 7.**
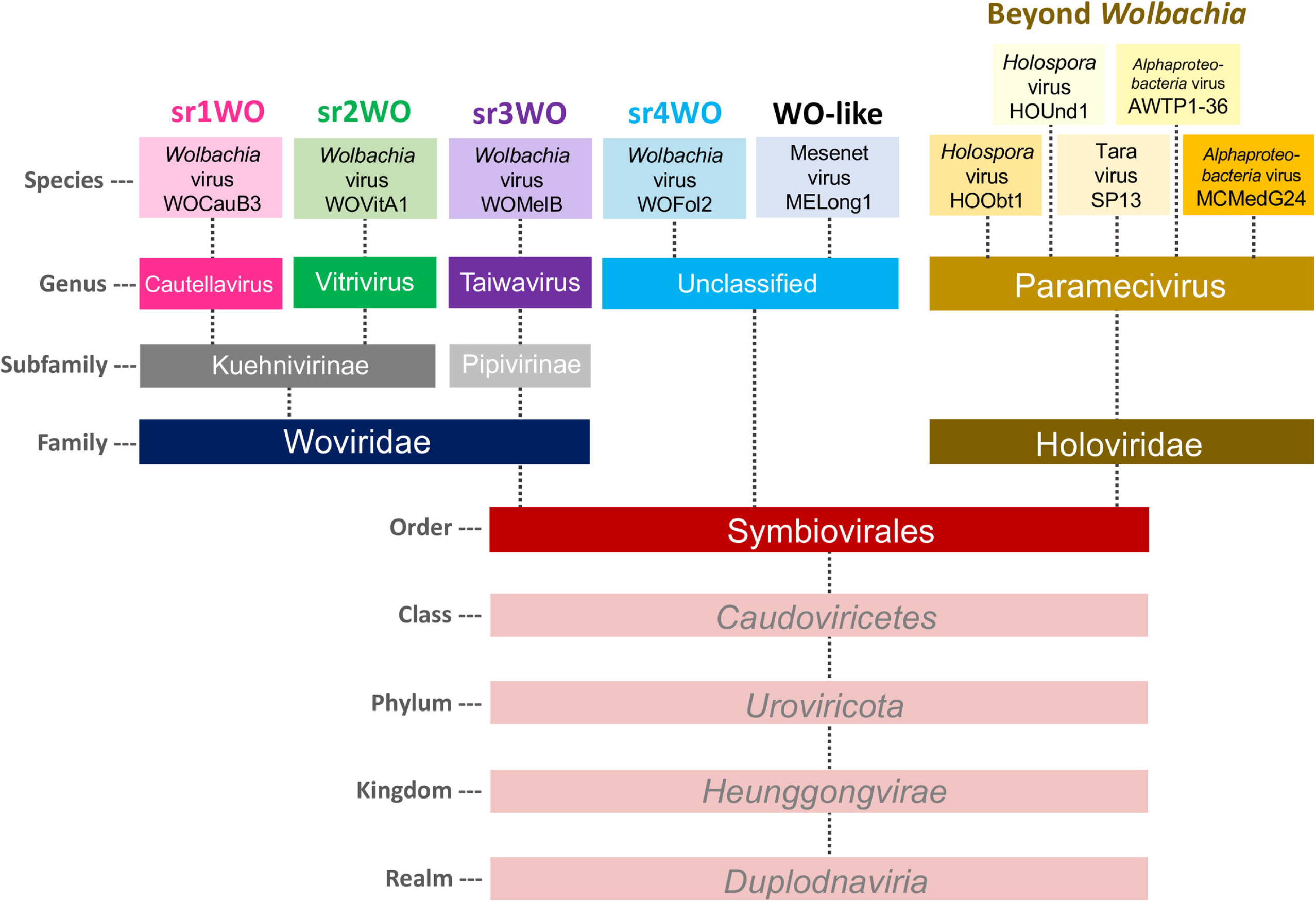
Comparative genomics supports a new order-level designation for prophage WO classification. Symbiovirales is proposed as a new taxonomic order of tailed phages within the class *Caudoviricetes*. It contains viruses that primarily infect *Wolbachia* (proposed family Woviridae) and other symbionts (proposed family Holoviridae). Two proposed subfamilies, Kuehnivirinae and Pipivirinae, distinguish the sr1WO/sr2WO and sr3WO clades (Figs 2 and 3, respectively). Three proposed genera of Woviridae include Cautellavirus (sr1WO), Vitrivirus (sr2WO), and Taiwavirus (sr3WO). sr4WO prophages are currently unclassified. Holoviridae contains a single proposed genus, Paramecivirus, that encompasses closely related prophages of *Holospora* and metagenome-assembled genomes (MAGs) from aquatic environments.

**Table 1.** The correlation between common name and proposed scientific name is listed for each phage WO exemplar variant and taxonomic rank.

| WO Exemplar Variant | Taxonomic Rank | Common Name | Proposed Scientific Name |
| --- | --- | --- | --- |
| WOCauB3 | Species | WOCauB3 | *Wolbachia* virus WOCauB3 |
|  | Genus | sr1WO | Cautellavirus |
|  | Subfamily | N/A | Kuehnivirinae |
|  | Family | Phage WO | Woviridae |
|  | Order | WO-like viruses | Symbiovirales |
| WOVitA1 | Species | WOVitA1 | *Wolbachia* virus WOVitA1 |
|  | Genus | sr2WO | Vitrivirus |
|  | Subfamily | N/A | Kuehnivirinae |
|  | Family | Phage WO | Woviridae |
|  | Order | WO-like viruses | Symbiovirales |
| WOMelB | Species | WOMelB | *Wolbachia* virus WOMelB |
|  | Genus | sr3WO | Taiwavirus |
|  | Subfamily | N/A | Pipivirinae |
|  | Family | Phage WO | Woviridae |
|  | Order | WO-like viruses | Symbiovirales |

We propose that all phage WO and WO-like viruses be classified in existing class *Caudoviricetes* (phylum *Uroviricota;* kingdom *Heunggongvirae;* realm *Duplodnaviria*) for tailed phages based on the presence of a tail module and observed tail-like structure in electron microscopy [34, 78]. We propose the new order Symbiovirales to recognize the association of these viruses with endosymbionts. Two proposed families, Woviridae and Holoviridae, are named after the first bacterial host identified for each family (*Wolbachia* endosymbionts of arthropods and *Holospora* endonuclear symbionts of *Paramecium*, respectively). Modules shared across the proposed Symbiovirales order are recombinase, replication, head, connector/baseplate, tail fiber, tail, and a putative lytic cassette (See Fig 8 for a summary of taxonomic traits).

**Fig 8.**
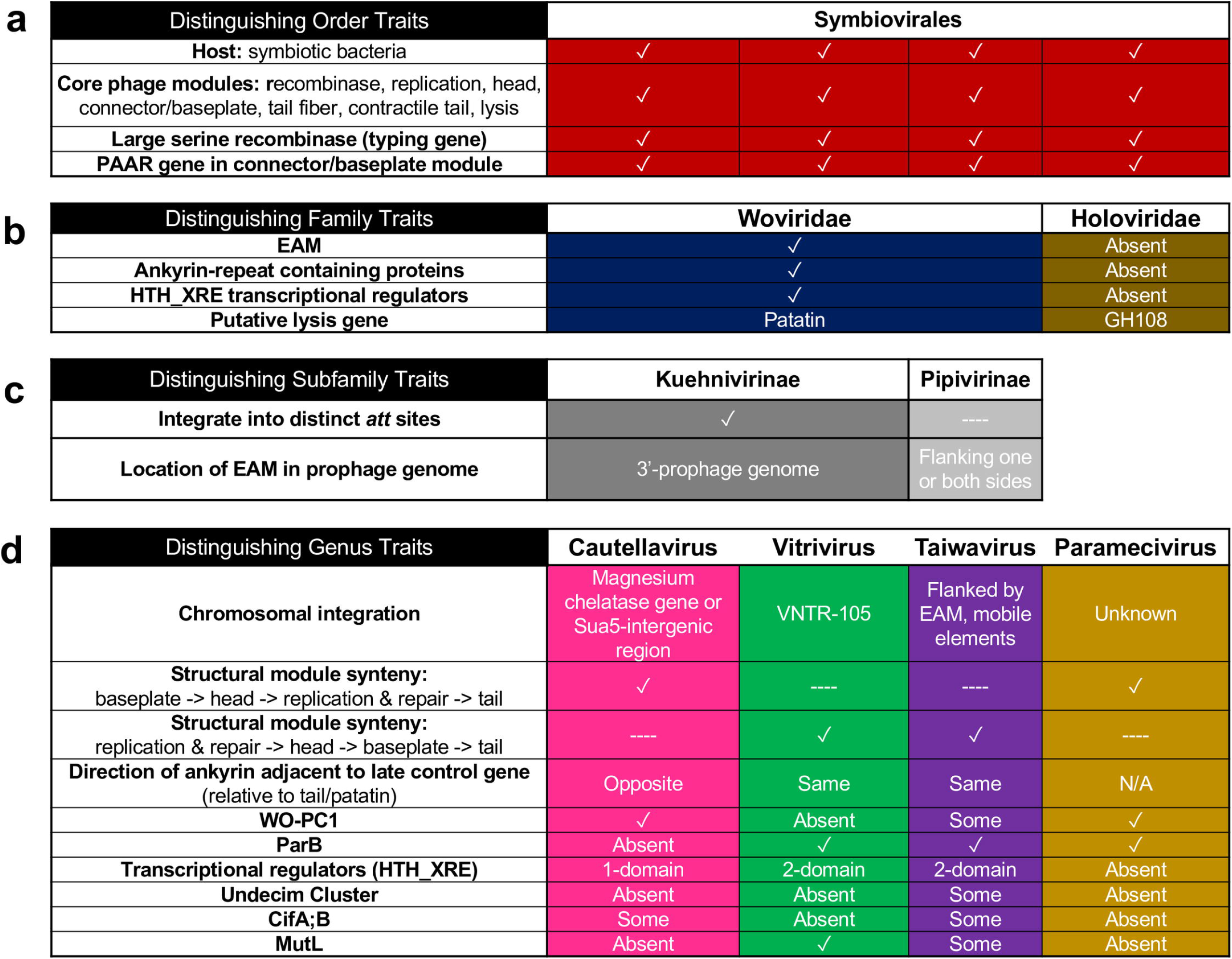
Linnaean classification of prophage WO-like viruses is supported by taxonomic traits at the order, family, subfamily, and genus level. (a) Proposed order Symbiovirales encompasses viruses that infect symbiotic bacteria, contain a large serine recombinase for integration and a PAAR gene in the connector/baseplate module, and feature a conserved set of core phage modules. They share nucleotide homology to *Wolbachia*’s prophages. (b) Subfamilies are classified by presence (Woviridae) or absence (Holoviridae) of an EAM and ankyrin repeat containing proteins. Woviridae may utilize patatin for lysis whereas Holoviridae encode a canonical GH108 endolysin. (c) Two proposed subfamilies address the diversity of chromosomal integration patterns and EAM location of prophages within the Woviridae family. (d) Proposed genera are further distinguished by multiple factors including structural module synteny, HTH_XRE domains, and genome composition.

The suggested family Woviridae encompasses all phage WO and prophage WO variants and is distinguishable by the presence of EAM and eukaryotic-like genes, a patatin-like phospholipase, and multiple ankyrin repeat containing proteins (Fig 8). Upon ICTV approval, Woviridae will be split into two subfamilies - Kuehnivirinae and Pipivirinae - named after the first purification of phage WO particles from *Ephestia kuehniella* [37] and *Culex pipiens* [35], respectively.

The proposed Kuehnivirinae will encompass two genera for phages that integrate into discrete *att* sites and feature 3’-placement of the prophage EAM. The first suggested genus of this subfamily, Cautellavirus, recognizes the sequenced genomes from *w*CauB phages [37, 38] and encompasses all sr1WO prophages (Fig 7). Cautellavirus core module synteny (replication, head, connector/baseplate) is inverted relative to other members of the proposed Woviridae; the ankyrin located between the tail module and putative lytic cassette is encoded on the opposite strand; and the genome does not contain a methylase/ParB protein (S1 Fig). The second suggested genus of this subfamily, Vitrivirus, recognizes the first fully sequenced genome from phage WOVitA1 particles [39] and encompasses all sr2WO prophages (Fig 7). Members of this genus feature discrete integration into the VNTR-105 locus, and the recombinase is adjacent to ankyrin repeats rather than WO-PC1.

Members of the proposed subfamily Pipivirinae are currently not associated with distinct *att* sites and are often flanked by EAM-like genes and transposases (S4 Table) on both ends of the integrated genome. Pipivirinae contains only one genus, Taiwavirus, named after the first prophage WO sequence fragment from *w*Tai [3, 78]. The proposed genus Taiwavirus will encompasses all sr3WO prophages (Fig 7) and is the most speciose genus of Symbiovirales. Likewise, it also features the greatest number of degraded prophage regions both within and across diverse *Wolbachia*. As more prophages are sequenced, it may be prudent to further classify this clade into subgenera based on presence or absence of the Undecim Cluster (Fig 2).

Finally, the WO-like prophages of *Candidatus* Mesenet longicola are likely classified as Woviridae due to nucleotide homology of structural genes and the presence of *cifA;B* containing EAM, but complete sequence information (specifically the recombinase and 5’-region beyond the CI loci) is necessary to definitively classify these phages. Likewise, the *w*Fol prophages will remain as *Unclassified* Woviridae until more genomes are sequenced to provide definitive taxonomic characteristics for the sr4WO variants. As more prophage WO genomes are sequenced, we propose using the srWO designation as a “common name” that roughly correlates with genus-level demarcation and referencing srWO when proposing future additions to the Woviridae taxonomy.

The proposed family Holoviridae includes the WO-like prophages from most non-*Wolbachia* metagenomic sequences and is currently comprised of phages from aquatic endosymbionts. They lack an EAM and ankyrin repeat containing proteins, feature a GH108 hydrolase rather than patatin-like phospholipase in the putative lytic cassette, and encode LexA and YqaJ that are generally absent from *Woviridae* genomes (Fig 6). Due to gene synteny and sequence homology of these prophage genomes, all species are currently classified into a single Paramecivirus genus. The first representatives of this genus were identified in *Holospora* spp., endonuclear symbionts of *Paramecium caudatum* and *P. bursaria* [97].

In summary, we propose that viruses should be classified as Symbiovirales based on reciprocal BLAST homology and shared gene content with core phage WO. The large serine recombinase can be used as a typing tool (Fig 3a) and intact genomes for inclusion should include (i) recombinase, (ii) replication and repair, (iii) connector/baseplate, (iv) tail fiber, (v) tail, and (vi) lytic modules. Woviridae are delineated by the presence of a eukaryotic association module (EAM), multiple ankyrin repeats, and a patatin-containing lytic module. Holoviridae are characterized by the absence of an EAM, lack of ankyrin repeats, and a GH108-containing lytic module.

## Discussion

The survey of 150 genomes coupled with manual annotations and comparative sequence analyses offers the most comprehensive overview of *Wolbachia* prophage WO genomics, distribution, and classification to date. From these analyses, we propose four major prophage WO variants corresponding with genus-level Linnaean taxonomy and support the creation of a new order Symbiovirales (within the *Caudoviricetes*) containing two distinct families, Woviridae and Holoviridae. Results presented above suggest that tailed, intact prophage WO genomes serve as a proxy for estimating prophage autonomy vs. domestication in the *Wolbachia* genome where multiple “degraded” prophages and WO-like Islands are indicative of prophage WO domestication by the bacterial host. WO regions enriched with transposable elements contribute to genome plasticity of the bacterial chromosome and may play a role in the domestication of these prophages. One such region, Octomom, has a putative WO origin in which a former EAM region is dynamically replicated or eliminated, and is associated with pathogenicity when not in a 1:1 ratio with its ancestral prophage. Finally, while there is currently no transformation system for *Wolbachia*, future applications may take advantage of conserved integration loci associated with each srWO and utilize the serine recombinase to introduce new genetic material into the bacterial chromosome.

### Establishment of the prophage WO database

To assist future analyses of prophage WO, a database of genomes discussed in this study is publicly available at https://lab.vanderbilt.edu/bordenstein/phage-wo/. The Prophage WO Database features sequence data, enhanced annotations, and phylogenetic tools to support: (i) identification of prophage WO regions in newly assembled *Wolbachia* genomes; (ii) annotation of the Undecim Cluster, cytoplasmic incompatibility (*cif*) genes, putative EAM genes, WO-PC2, and other WO-associated regions; and (iii) taxonomic classification of prophage WO-like viruses.

## Methods

### Prophage WO genome maps and chromosomal integration patterns

Prophage WO regions were manually retrieved from sequenced *Wolbachia* genomes in GenBank via BLASTN searches against each individual *Wolbachia* genome in the Nucleotide (NR/NT) and WGS databases [43]. Genomes from WOCauB3, WOVitA1, WOMelB, WOPip5, and WOFol3 were the primary reference genomes used for each search. Because most prophage regions were incomplete and located at the ends of contigs, we selected more complete assemblies for comparative genomics: *w*Ri, *w*Ana, *w*Suzi, *w*VitA, *w*Ha, *w*Mel, *w*Pip, *w*No, *w*Au, *w*Irr, *w*Fol, *w*AlbB, *w*Mau, and the previously described prophage genomes WOKue, WOCauB2, WOCauB3, WOSol, WORecA, and WORecB (See S1 Table for accession numbers). All genomes were reannotated in Geneious Prime v2019.2 using the InterProScan [98] plug-in along with information from BLASTP [99], Pfam [100], HHPRED [101], ISFinder [102], and SMART [103] databases. Prophages were then organized into groups based on similar gene content and module organization. Whole genome alignments were performed with the Mauve [104] plug-in in Geneious.

Prophage genomic boundaries for sr1WO and sr2WO were defined by 5’ and 3’ homology to a known *attP* site (discussed below). Prophage genomic boundaries for sr3WO and sr4WO were identified by translating each prophage gene and “walking out” from the structural modules by using a BLASTP of each gene product against the core *Wolbachia* genome. If a gene was identified in most *Wolbachia* strains, including those infecting nematodes, as well as in the closely related genera *Ehrlichia* and *Anaplasma*, it was considered a core *Wolbachia* gene and not included in the prophage annotation. If a gene was only present in WO-like regions of other *Wolbachia* genomes, it was considered a phage-associated gene. Because the HTH_XRE transcriptional regulators (WO-PC2) were identified in phage purifications from WOCauB3 and WOVitA1, any genes located between the structural modules and WO-PC2 were considered part of the prophage genome. Through this method, we identified flanking 5’ and 3’ transposases that separated phage-associated genes and the bacterial chromosome in sr3WO and sr4WO regions. Because some transposable elements did not fall within the known IS Groups for *Wolbachia* [2], they were comparably annotated to IS Family using ISFinder.

Chromosomal integration patterns were analyzed by similarly aligning all circular genomes based on the putative origin of replication, *ori* [61]: WD1027 (CBS domain-containing)-like genes were oriented in the reverse direction and WD1028 (*hemE*)-like genes were oriented in the forward direction. The nt-distance from *ori* to the prophage recombinase, or 5’-gene, was divided by the length of the total *Wolbachia* genome and multiplied by 100 for a % distance from *ori*. The *w*VitA and *w*Rec genome arrangements may not be exact as they contain multiple scaffold breaks and genome orientation was estimated based on homology to closely related genomes.

### Recombinase homology and phylogenetics

Large serine recombinase genes from each reference genome were translated and aligned using the MUSCLE [105] plugin in Geneious. The best model of evolution, according to corrected Akaike information criteria, was determined by ProtTest [106, 107] and the phylogenetic tree was constructed using default parameters of the MrBayes [108] plugin in Geneious with Rate Matrix=jones and Rate Variation=invgamma. A Consensus Tree was built with a support threshold of 50% and burn-in of 10%.

### Phage WO *att* sites

The *attP* sites for WOVitA1 and WOCauB3 were previously identified by sequencing active phage particles and confirmed with PCR and Sanger sequencing [39]. Each *attP* sequence was submitted as a BLASTN query against *Wolbachia* genomes harboring similar prophage haplotypes to identify specific *attL* and *attR* sites. The *attB* sites were predicted by concatenating chromosomal sequences adjacent to *attL* and *attR.* The predicted *attB* sites were then used as queries in a BLASTN search against *Wolbachia* genomes to confirm that the sequences exist, uninterrupted, in chromosomes lacking similar prophage haplotypes.

### Phage WO beyond *Wolbachia*

Contigs containing WO-like prophage regions in *Holospora*, *Orientia, Candidatus* Mesenet, and multiple metagenome-associated taxa were identified by a BLASTP query of prophage WO sequences against the NCBI database. The nucleotide sequence for each homolog (usually a contig in the WGS database) was manually inspected for WO-like regions. If detected, the boundaries of each prophage region were determined using the similar “walk out” BLASTP approach described above, looking for homology to other phage or bacterial genes. All non-Anaplasmataceae prophage genomes had concise boundaries (recombinase and lysis module) that did not include an EAM.

### Identification of Gene Transfer Agents

The genome annotations used for comparative genomics were manually inspected for keywords *phage, capsid*, and *tail*. Any gene not within an annotated prophage WO region was translated and a BLASTP was performed against the NCBI database. Based on top hits, genes were binned into “WO-like” indicating homology to phage WO and “GTA” indicating homology to HK97 phage.

### Taxonomic Classification

The proposed taxonomic classification of phage WO was drafted in accordance with ICTV guidelines for genome-based taxonomy [109] and will be formally reviewed by the Committee in the next cycle. Specifically, it is recommended that phages should be assigned the same species if their genomes are more than 95% identical; assigned the same genus if genomes share 80% nucleotide identity across the genome length and form monophyletic groups based on a phylogenetic tree of signature gene(s); assigned the same subfamily (optional) if they share a low degree of sequence similarity and the genera form a clade in a marker tree phylogeny; assigned the same family if they share orthologous genes and form a cohesive and monophyletic group in a proteome-based clustering tool; and assigned the same order when two or more families are related. Prophage WO taxonomic classification satisfied all demarcation criteria except for genus designation. At the genus level, due to the high variability of the EAM, we applied alternative criteria: genomes should (i) share >70% nucleotide homology across >30% of the genome; (ii) form a distinct phylogenetic clade based on the amino acid sequence of the signature typing gene, large serine recombinase; and (iii) demonstrate shared gene and module synteny.

## Supplementary Figures

**S1 Fig. Cautellavirus (sr1WO) genome maps.** Genome maps of sr1WO prophage regions where genes are drawn to scale in forward and reverse directions. Predicted physical structures are illustrated to the left of each genome. All genomes contain tail modules with the exception of the partial WOVitA2 sequence. Prophage WO Core Genes are shaded in blue and predicted EAM genes are shaded in gray. Genes of similar function are similarly color-coded according to the figure legend. Locus tags, if available, are listed in italics above the genes. The large, black diagonal lines between the recombinase and transposase in WORiC and WOSuziC represent post-integration rearrangement of the prophage region in the *Wolbachia* chromosome. Dashed lines represent breaks in the assembly whereas small diagonal lines represent a continuation of the genome onto the next line. Arrows with diagonal stripes represent genes that may be pseudogenized relative to homologs in other prophage WO genomes. The putative function for each structural gene is discussed in S1 Text.

**S2 Fig. Vitrivirus (sr2WO) genome maps.** Genome maps of sr2WO prophage regions where genes are drawn to scale in forward and reverse directions. Predicted physical structures are illustrated to the left of each genome. WOVitA1-like prophage genomes encode all structural modules (shaded in blue) and an EAM (shaded in gray) whereas WORiA-like prophage genomes encode an intact head module, recombinase, lysozyme, AAA16, and disrupted connector. They lack most other modules. Genes of similar function are similarly color-coded according to the figure legend. Locus tags, if available, are listed in italics above the genes. Dashed lines represent breaks in the assembly whereas small diagonal lines represent a continuation of the genome onto the next line. Arrows with diagonal stripes represent genes that may be pseudogenized relative to homologs in other prophage WO genomes. The putative function for each structural gene is discussed in S1 Text.

**S3 Fig. Taiwavirus (sr3WO) genome maps.** Genome maps of sr3WO prophage regions where genes are drawn to scale in forward and reverse directions. Three *w*Pip prophages exist as one contiguous prophage region in the *Wolbachia* genome and are illustrated here as WOPip1, WOPip2, and WOPip3 (based on [110]). Predicted physical structures are illustrated to the left of each genome. Prophage WO Core Genes are shaded in blue and predicted EAM genes are shaded in gray. Genes of similar function are similarly color-coded according to the figure legend. sr3WO is comprised of highly variable genomes that are often flanked by mobile elements (transposases are shown in yellow). They generally contain a recombinase, connector/baseplate, head, and EAM with only a few genomes encoding a complete tail. Prophages in this group often contain *cifA;B* (pink). Locus tags, if available, are listed in italics above the genes. Dashed lines represent breaks in the assembly whereas small diagonal lines represent a continuation of the genome onto the next line. Arrows with diagonal stripes represent genes that may be pseudogenized relative to homologs in other prophage WO genomes. The putative function for each structural gene is discussed in S1 Text.

**S4 Fig. Taiwavirus (sr3WO and sr3WO-Undecim Cluster) genome maps.** Genome maps of sr3WO prophage regions where genes are drawn to scale in forward and reverse directions. WOIrr is one contiguous prophage region in the *Wolbachia* genome that is illustrated here as Segment 1 and Segment 2. A subset of sr3WO prophages is further categorized by the presence of a highly conserved WD0611-WD0621 like region, termed the Undecim Cluster (navy blue). Predicted physical structures are illustrated to the left of each genome. Prophage WO Core Genes are shaded in blue and predicted EAM genes are shaded in gray. Genes of similar function are similarly color-coded according to the figure legend. sr3WO is comprised of highly variable genomes that are often flanked by mobile elements (transposases are shown in yellow). Prophages in this group often contain *cifA;B* (pink). Locus tags, if available, are listed in italics above the genes. Dashed lines represent breaks in the assembly whereas small diagonal lines represent a continuation of the genome onto the next line. Arrows with diagonal stripes represent genes that may be pseudogenized relative to homologs in other prophage WO genomes. The putative function for each structural gene is discussed in S1 Text.

**S5 Fig. Taiwavirus (sr3WO-Undecim Cluster) genome maps.** Genome maps of sr3WO prophage regions where genes are drawn to scale in forward and reverse directions. This subset of sr3WO prophages is further categorized by the presence of a highly conserved WD0611-WD0621 like region, termed the Undecim Cluster (navy blue). Predicted physical structures are illustrated to the left of each genome. Prophage WO Core Genes are shaded in blue and predicted EAM genes are shaded in gray. Genes of similar function are similarly color-coded according to the figure legend. sr3WO is comprised of highly variable genomes that are often flanked by mobile elements (transposases are shown in yellow). Prophages in this group often contain *cifA;B* (pink). Locus tags, if available, are listed in italics above the genes. Dashed lines represent breaks in the assembly whereas small diagonal lines represent a continuation of the genome onto the next line. Arrows with diagonal stripes represent genes that may be pseudogenized relative to homologs in other prophage WO genomes. The putative function for each structural gene is discussed in S1 Text.

**S6 Fig. Unclassified (sr4WO) genome maps.** Genome maps of sr4WO prophage regions where genes are drawn to scale in forward and reverse directions. To date, sr4WO prophages have only been identified in the parthenogenic strain of *Folsomia candida*, *w*Fol. WOFol2 is one contiguous prophage region in the *Wolbachia* genome that is illustrated here as Segment 1 and Segment 2. Likewise, the WOFol3 prophage region is illustrated as three segments. Predicted physical structures are illustrated to the left of each genome. Prophage WO Core Genes are shaded in blue and predicted EAM genes are shaded in gray. Genes of similar function are similarly color-coded according to the figure legend. Locus tags, if available, are listed in italics above the genes. Small diagonal lines represent a continuation of the genome onto the next line. Arrows with diagonal stripes represent genes that may be pseudogenized relative to homologs in other prophage WO genomes. The putative function for each structural gene is discussed in S1 Text.

**S7 Fig. WO-like Island genome maps.** Genome maps of WO-like Islands where genes are drawn to scale in forward and reverse directions. These regions contain only one structural module and/or group of WO-related genes. Regions flanked by assembly breaks (i.e., WORecB, WORecA, and *w*VitA) are tentatively classified as WO-like Islands due to lack of a full-length prophage in the genome assembly. Names are based on the original author’s description. If it was identified as a prophage in the genome announcement, the reported WO name is listed here. Otherwise, the name simply refers to the encoding *Wolbachia* genome. Many WO-like Islands contain *cifA;B*; some Islands (i.e., *w*No, *w*VitA, WOMau4, and WOAlbB3) contain both Type III *cifA;B* (pink) and the Undecim Cluster (navy blue). Predicted physical structures are illustrated to the left of each genome. Prophage WO Core Genes are shaded in blue and predicted EAM genes are shaded in gray. Genes of similar function are similarly color-coded according to the figure legend. Locus tags, if available, are listed in italics above the genes. Dashed lines represent breaks in the assembly. Arrows with diagonal stripes represent genes that may be pseudogenized relative to homologs in other prophage WO genomes. The putative function for each structural gene is discussed in S1 Text.

**S8 Fig. *In silico* predictions of phage WO attachment (*att*) sites.** An integrated prophage sequence contains left and right attachment sites (*attL* and *attR*, respectively) at the points of chromosomal integration. Half of the *att* site is phage-derived (green); the other half is bacterial derived (black). If the DNA sequence of the bacterial attachment site (*attB*, black) is known, a nucleotide alignment of the intact sequence with the integrated prophage genome will correlate with 5’-(*attL*) and 3’-(*attR*) prophage boundaries. (a) WORiC, a member of sr1WO, integrates into *w*Ri’s magnesium chelatase gene. By aligning an intact copy of this gene (WD0721) from closely related *w*Mel that does not harbor sr1WO, (b) the juncture points of the disrupted magnesium chelatase indicate the *attL* and *attR* sites for the WORiC prophage region within the *w*Ri genome. (b) The phage attachment site (*attP*, green) is predicted *in silico* by concatenating the non-*Wolbachia* portions of the *attL* and *attR* sites. (c) Likewise, this method can also be applied when the bacterial integration locus is intergenic. The homologous intergenic region of closely related, sr1WO-free *w*Pip can be used to predict *att* sites for WOCauB3. Nucleotides in orange represent a common region, O, that is shared by all four *att* sites. This method was adapted from [39] where the *attP* site was used to predict the *attB* site of sr2WO phages.

**S9 Fig. RT is associated with duplication, inversion, and recombination of the prophage WO genome.** (a) The WOMelB prophage genomes of *w*Mel2_a and *w*Mel2_b are duplicated relative to the *w*Mel reference genome [72]. (b) The entire WORiB prophage region is duplicated in *w*Ri [19]. (c) WOHa1 encodes a second, pseudogenized *cifA;B*-containing region relative to closely related WOAuA, WORiB, WOSuziB, and WOSol prophages. (d) A ligase-containing region is duplicated in *w*Fol’s WOFol1 and WOFol2 [56]. (e) Based on homology to other prophage regions (Fig 2), the connector/baseplate should be adjacent to a head module and the WOPC-2 and replication genes should be oriented in the opposite direction; this indicates a likely insertion and/or recombination in the WOFol3 prophage region. (f) The WOIrr head module is inverted relative to other sr3WOs. Genes are illustrated as arrows; putative gene annotations are labeled in S1-S7 Figs. In each example, the regions of chromosomal rearrangement are highlighted in light orange and flanked by at least one RT.

**S10 Fig. Comparative genomics of Octomom-like variants across diverse *Wolbachia*.** Octomom (yellow-orange) and Octomom-like (green) regions are illustrated for *w*MelCS, *w*Mel, *w*SYT clade, and *w*Pip. Characteristics of each region are listed next to the genome schematic. Notably, the wMelCS genome, representative of the dynamic wMelPop, is distinguished from other variants by intact, flanking reverse transcriptases of group II intron origin (RT) on both sides. wPip, the only *Wolbachia* Supergroup B variant, is the most divergent and not associated with an RT, MutL or ankyrin repeat. Rather it is adjacent to WP1349, another gene that has been horizontally transferred between phage and arthropod [71].

**S11 Fig. Prophage WO encodes a putative lytic cassette.** Adjacent to the tail module of most prophage WO variants are three phage lysis candidates: ankyrin repeat containing protein (not shown), holin-like, and patatin-like phospholipase. (a) Similar to canonical holins, the prophage WO gene product encodes a single N-terminal transmembrane domain with no predicted charge. It is smaller than 150 amino acid residues, features a C-terminal coiled coil motif, and has a highly charged C-terminal domain. Unlike canonical holins, however, it is adjacent to a patatin-like gene rather than a characterized endolysin. (b) The prophage WO holin-like peptide shares 41.1% amino acid identity to a homolog in the non-*Wolbachia* prophage from the Tara Oceans Project that is directly adjacent to a GH108 lysozyme (complete genome illustrated in Fig 6). (c) A Mauve alignment of these genomic regions (core phage modules only; EAM not included) indicates 50.3% nucleotide identity across the majority of the sequence, including the holin-like gene (marked with a gold star). The similarity of these prophages suggest that prophage WO may utilize a similar holin-like gene with a different lytic enzyme (i.e., patatin rather than lysozyme) to lyse the bacterial cell.

**S12 Fig. *Wolbachia* contains both prophage regions and GTA-like genes scattered through the chromosome.** (a) Circular *w*Mel contains three prophage WO-like regions (teal) and multiple genes with homology to GTAs (orange) scattered throughout the genome, illustrated relative to the putative origin of replication (*ori*, gray). The Undecim Cluster is highlighted in navy blue, *cifA;B* are highlighted in pink, and *wmk* is highlighted in purple. (b) GTAs are present in at least one strain of each *Wolbachia* Supergroup except Supergroups J and L. They are also present in closely related Anaplasmataceae genera.

**S13 Fig. Distance matrices of GTA nucleotide homology indicate evolution with the *Wolbachia* chromosome.** Nucleotide alignments of GTA genes (a) portal, (b) BRO599, (c) TIM barrel, (d) major capsid, (e) head-tail connector, and (f) terminase indicate strict delineation based on *Wolbachia* supergroup. This supports evolution with the *Wolbachia* chromosome rather than independent evolution of a phage genome.

## Supplementary Tables

**S1 Table. Prophage WO genes are associated with arthropod-infecting *Wolbachia*.** *Wolbachia* genomes are listed according to (A) host phylum; (B) *Wolbachia* supergroup; (C) *Wolbachia* name (D) host species and (E) host strain/lineage, if applicable; (F) NCBI accession number; (G) genome assembly status; (H) identification of prophage WO core genes; (I) identification of CI genes; and (J) identification of the Undecim Cluster. *Wolbachia* strains that did not include official names in the assembly reports are listed here using a capital letter for host genus and two to three lowercase letters for host species. “Highly pseudogenized” in column H indicates that the prophage genome is highly pseudogenized and encodes very few Core WO genes. (*) indicates that the genome lacks a complete Undecim Cluster but encodes WD0616 and/or WD0621 homologs. (**) indicates that the genome was not included as Source Data for Fig 1b due to incomplete genome information.

**S2 Table. *w*Mhie encodes prophage WO genes.** *w*Mhie, a *Wolbachia* endosymbiont from the nematode *Madathamugadia hiepei*, encodes four genes that are conserved throughout phage WO’s transcriptional regulation and replication/repair modules. Each gene is listed by locus tag, annotation, and nucleotide homology to prophage WOVitA1.

**S3 Table. cifA and *cifB* genes are associated with *Wolbachia* Supergroups F and T.** *cifA* and *cifB* are identified in Supergroups F and T. NCBI accession numbers and genomic coordinates (or locus tags) are provided for each locus.

**S4 Table. Diversity of prophage WO mobile elements.** All mobile elements, both flanking and internal, are listed for each prophage WO genome according to original genome annotations and ISFinder [102]. The sr1WO group and WOVitA1-like prophages of the sr2WO group do not feature transposases on the 5’- and 3’-flanking regions. The WORiA-like prophages of the sr2WO group are associated with 3’-transposases; these correlate with putative truncations of the prophage regions. Most genomes within the sr3WO group feature mobile elements on both 5’- and 3’-ends. IS refers to Insertion Sequence Family; RT refers to reverse transcriptase of group II intron origin; Rpn refers to recombination promoting nuclease. (*) indicates a sequencing gap or artificial join in the *Wolbachia* genome. Complete sequence information is unknown. (**) indicates that these prophage sequences were obtained from contigs and may be segmented in the *Wolbachia* chromosome; the exact 5’ and 3’ ends are uncertain. Genomic locations for each mobile element are illustrated in S1-S7 Figs.

**S5 Table. *Wolbachia* GTA genes.** The annotation of *Wolbachia*’s distributed GTA genes is based on a BLASTP against NCBI Conserved Domains; E-values are listed in column B.

## Supplementary Text

**S1 Text: Phage WO Structural Modules.**

Phage WO structural genes are organized into head, connector/baseplate, tail, and tail fiber modules. The predicted function of each gene is discussed based on conserved protein domains and homology to other model systems, including lambda, T4, P2, and Mu phages.

## Acknowledgements

We would like to thank Evelien Adriaenssens for helpful guidance with the taxonomic classification of prophage regions.

## Author Contributions

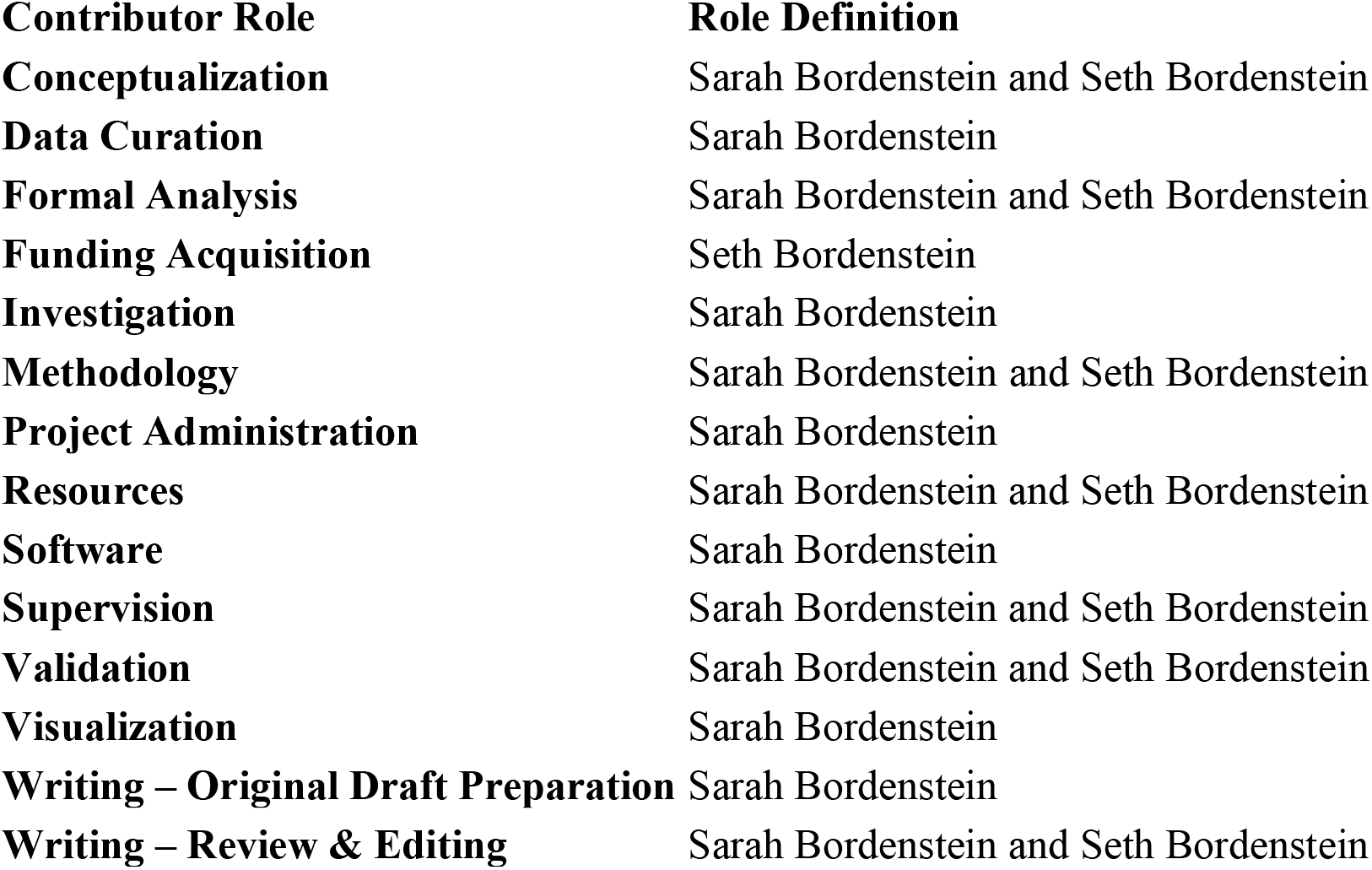

## List of Abbreviations

CI: cytoplasmic incompatibility
EAM: eukaryotic association module
GTA: gene transfer agent
ICTV: International Committee on Taxonomy of Viruses
IS: insertion sequence
HTH: helix-turn-helix
NCBI: National Center for Biotechnology Information
Rpn: recombination-promotion nuclease
RT: reverse transcriptase of group II intron origin
VNTR: variable number tandem repeat
WO-PC1: WO protein cluster 1
WO-PC2: WO protein cluster 2

## Notes

https://lab.vanderbilt.edu/bordenstein/phage-wo/

### Competing Interest Statement

The authors have declared no competing interest.

