## Supplementary material for "The most widespread phage in animals: Genomics and taxonomic classification of Phage WO": S1 Text

### S1 Text: Phage WO Structural Modules

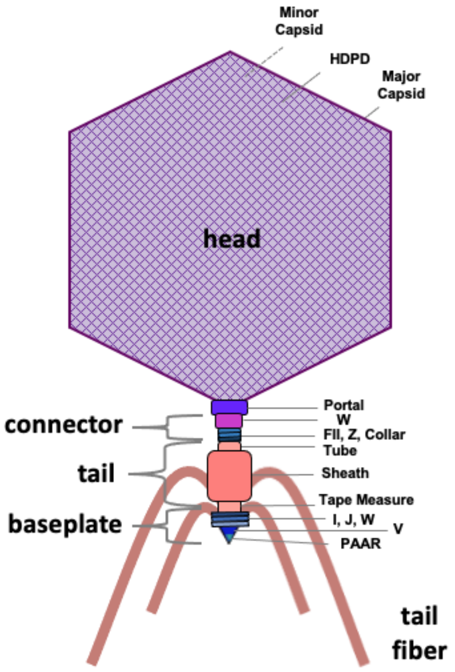


**Fig 1. Prophage WO encodes conserved structural proteins.** An illustration of phage WO's putative structural proteins, as determined by protein annotations using NCBI, SMART Protein, and HHpred.

Phages of obligate intracellular endosymbionts are less understood than phages of free-living bacteria due to an inability to efficiently purify and genetically manipulate these viruses. Therefore, we report *predicted functions* of phage WO structural genes based on conserved protein domains and homology to other model systems, including lambda, T4, P2, and Mu phages. Citations are provided for each annotation; the actual function of each gene is to be determined.

*Head*

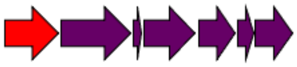

**Fig 2. Genome map of Phage WO head module.** From left to right: Ankyrin – Terminase – gpW – Portal – Minor capsid – HDPD – Major capsid

The gene cluster comprising the head module is highly conserved across all prophage WO variants. The **portal** is the initiator of head assembly and forms a structural hole enabling DNA passage during packaging and ejection [1]. It also serves as a connection point for the head and tail proteins. **Minor capsid C** (*orf7*), identified by its peptidase family S49 domain, creates an initiator structure upon which the **major capsid E** forms the phage procapsid [2]. *Orf7* is historically used to PCR-detect prophage WO [3, 4]. Finally**, head decoration protein D** stabilizes the structure, allowing for full-length packaging of the genome, and completes the mature phage head particle [5]. Phage DNA is translocated and packaged into the head by the **terminase** and then stabilized by **gpW** via plug formation or direct binding to the DNA [6]. Unique to *Wolbachia*'s prophage WO, and directly upstream from the portal, is a conserved **ankyrin repeat protein** with C-terminal transmembrane domains. Prophages without this complete ankyrin are typically associated with an incomplete head module or disruptions/truncations of essential genes within the region (i.e., WOVitA2, WOMelA, WOHa1). It is unknown, however, if this ankyrin is part of the complete head module.

*Connector/Baseplate*

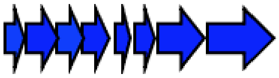

**Fig 3. Genome map of Phage WO connector/baseplate module.** From left to right: gpFII – gpZ – Collar – gpV – PAAR – gpW – gpJ - gpI

The connector/baseplate module is also associated with all prophage WO groups. Conserved protein domains match those of the well-described lambda, T4, and P2 bacteriophages. The first three genes likely comprise the head-tail connector, or neck, region. **Minor tail protein Z** (gpZ) is involved in the completion of bacteriophage lambda tail so that it can be attached to the head [7] and **gpFII** joins the phage head and tail in the final stage of virion assembly [8]. **Collar** shares structural homology with the ring structure of a contractile R-type bacteriocin (HHpred, 100% probability, E-value 1.7e-30) where it binds the tail tube to the contractile sheath [9]. In Mu-like phages, a similar protein (gp37; HHpred, 99.55% probability, E-value 7.2e-14) forms part of the neck, connecting the head and tail [10]. The final five genes are involved with baseplate formation and host adsorption. **gpJ** and **gpI** comprise baseplate wedges in bacteriophage P2 [11] while **gpW**, similar to gp25 of bacteriophage T4, forms the outer wedge of the baseplate and has been suggested to have lysozyme activity required for cell-entry [12]. **GpV** forms the P2 tail-spike protein and irreversibly punctures and adsorbs to bacterial host cells [13]. Finally, **PAAR** repeat-containing proteins have been annotated in numerous T4-like phages [14], and crystal structures of the Type VI secretion system show that they sharpen the tail-spike and selectively attach cytotoxic effector proteins [15, 16].

*Tail*

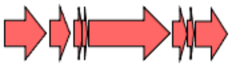

**Fig 4. Genome map of Phage WO tail module.** From left to right: Tail sheath – Tail tube – gpG/GT – Tape measure – gpU –gpX – Late control

The prophage WO tail module is comprised of eight conserved genes. Based on protein analysis, the tail likely utilizes a **contractile sheath** that surrounds the **tail tube** and contracts during bacterial infection to drive the tube through the outer membrane and deliver the viral genome [17]. **The tape measure protein** is located inside the tube and, as its name implies, determines the length of the phage tail [18]. **GpX** is a phage tail protein with unknown function. Studies in P2-like phages indicate that the protein shares sequence and structural homology to peptidoglycan-binding LysM domains [19], but this activity has not been confirmed for gpX. **GpU** and **late control** (gpD) share conserved protein domains with P2-like phages and are thought to be involved in tail assembly and late control, respectively. Lastly, we report a conserved translational frameshift between the tail tube and tape measure genes that encodes tail assembly chaperone genes **gpG** and **gpGT** [20]. Their sequential formation of a spiral structure is essential for proper tail tube polymerization [21]. The characteristic overlapping open reading frames (ORFs) between these tail genes has been identified in all tailed phage WO genomes along with either a 5’-TTTTTTG-3’ or 5’-TTTTTTT-3’ slippery sequence, correctly positioned to produce the characteristic -1 frameshift (Fig 5). BLASTp analysis places the corresponding WOVitA1 gp43 gene product in the Flu_Mu_gp41 family (E-value 1.41e-19), a well-described λ gpG/GT analog [22].

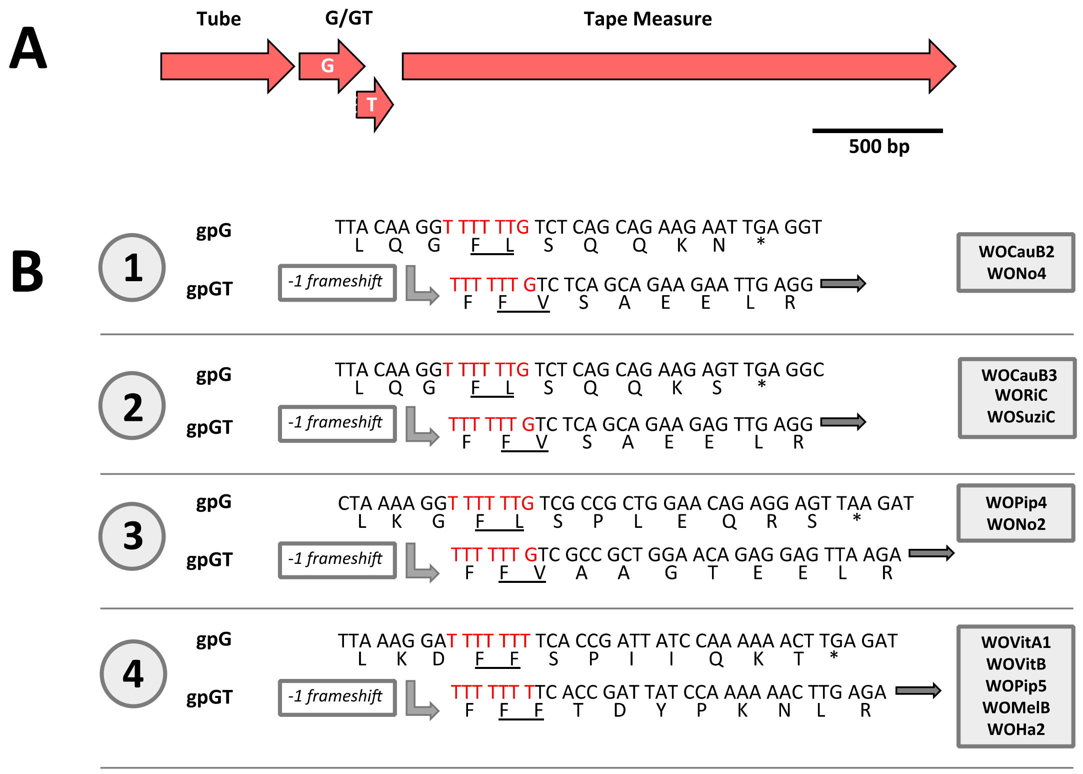


**Fig 5. Prophage WO genomes contain a conserved translational frameshift in the tail assembly chaperone genes, *gpG* and *gpGT*.** (a) The tail module of the prophage WO genome contains two overlapping ORF’s between the tube and tape measure genes. (b) A common slippery sequence of either 5'-TTTTTTG-3' or 5'-TTTTTTT-3' was identified in tailed prophage WO regions and is strongly suggestive of a -1 frameshift. The predicted "slippery sequence" is highlighted in red. The prophages are organized into groups that have identical sequences in the frameshift region.

##### *Tail fiber*

Phage WO’s putative tail fiber module consists of six highly conserved genes. Components of this module are historically annotated as hypothetical proteins due to a lack of sequence homology with well-studied model systems. Algorithms based on structural prediction rather than sequence homology, however, indicate that the module is a putative tail fiber network with gene synteny that closely resembles the receptor-binding proteins of *Listeria* phage A511. Here we discuss a putative model for the phage WO tail fiber based on cryo-electron microscopy, cryo-electron tomography, and X-ray crystallography of syntenic A511 homologs [23].

a

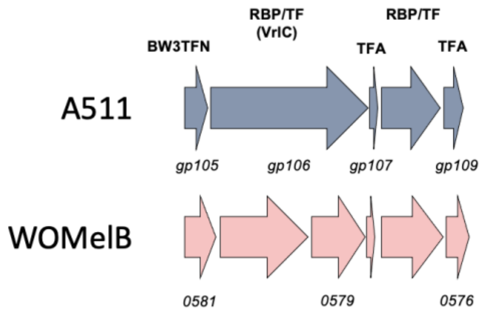


b

| WOMelB locus tag | HHpred | Probability (%) | E-value | % aa identity WOMelB : A511 (MUSCLE) | A511 locus tag | HHpred | Probability (%) | E-value | A511 Annotation (Guerrero-Ferreira et al., 2019) |
|---|---|---|---|---|---|---|---|---|---|
| **WD0581** | Gp105 | 100 | 1.50E-38 | 17.7% | **gp105** | Gp105 | 100 | 2.10E-46 | Baseplate Wedge 3 Tail Fiber Network (BW3TFN) |
| **WD0580** | DUF4815 | 100 | 5.10E-117 | 16.5% | **gp106-NTD** | DUF4815 | 100 | 1.10E-108 | Receptor-Binding Protein (RBP) or Tail Fiber (TF), previously VrlC |
| **WD0579** | Lectin-like | 97.39 | 0.0068 | 11.2% | **gp106-CTD** | Lectin-like | 96.05 | 0.23 | |
| **WD0578** | DUF2730 | 84.93 | 10 | 17.6% | **gp107** | Prefoldin | 94.77 | 0.81 | Tail Fiber Assembly chaperone (TFA), putative |
| **WD0577** | Factor H-binding protein (FHbp) | 78.83 | 160 | 14.1% | **gp108** | Tail fiber | 99.88 | 4.30E-18 | Receptor-Binding Protein (RBP) or Tail Fiber (TF) |
| **WD0576** | Tail fiber assembly protein | 92.7 | 4.3 | 16.7% | **gp109** | Phage-like element PBSX | 99.9 | 8.20E-23 | Tail Fiber Assembly chaperone (TFA), putative |

**Fig 6. Phage WO encodes a putative tail fiber module based on gene synteny and comparable structural annotation with *Listeria* phage A511**. (a) Shared gene synteny of A511 and WOMelB**.** The DUF4815 and lectin-like domains, formerly annotated as VrlC, are encoded by separate genes in WOMelB and fused in A511. (b) HHpred-based predictions are highly similar for WOMelB and A511, but the phage proteins do not share amino acid homology. WOMelB predictions are highlighted in pink and A511 predictions are highlighted in gray. Side-by-side comparisons suggest similar function for syntenic genes in the module.

**WD0581** shares structural homology to a baseplate wedge 3 tail fiber network protein, *BW3TFN* (HHpred, 100% probability, E-value 1.5e-38) of phage A511. Like its T4 ortholog, gp8, the A511_Gp105 protein forms a dimer that is either involved in tail fiber attachment or serves as the integral component of the fiber network [23].

**WD0580** and **WD0579**, initially annotated as virulence factors VrlC.1 and VrlC.2 based on weak amino acid homology to a pathogenicity island of the sheep pathogen *Dichelobacter nodosus [24]*, are predicted to encode the same domains as A511_Gp106. WD0580 and the A511_Gp106-NTD share significant structural homology to DUF4815 (WD0582: HHpred, 100% probability, E-value 5.1e-117) while WD0579 and A511_Gp106-CTD share slightly weaker homology to a lectin-like domain (WD0583: HHpred, 97.39% probability, E-value 0.0068). Structural assays of the A511 homolog identify Gp106 as a receptor-binding protein (*RBP*) or tail fiber component (*TF*) that forms a pyramid structure at the proximal part of the tail fiber. The pyramid initially points towards the phage capsid but re-orients towards the host to facilitate binding of the phage particle to the cell wall upon tail sheath contraction [23].

The role of A511_Gp107 in phage biology is less clear, but it is thought to be involved in the pyramid structure and possibly interact with the cell wall [23]. The phage WO counterpart in the gene synteny, **WD0578**, is similar to a phage-associated Domain of Unknown Function protein (DUF2730; HHpred, 84.93% probability, E-value 10). While its function is unknown, module conservation across prophage WO variants strongly suggests a role in tail fiber assembly.

**WD0577** shares weak structural homology to Factor H-binding protein (FHbp; HHpred, 78.83% probability, E-value 160), a virulence protein of *Neisseria meningitidis*, and extracellular matrix glycoprotein Laminin subunit beta (HHpred, 78.28% probability, E-value 97). Both proteins are involved in cell binding of eukaryotic cells. Likewise, syntenic A511_gp108 forms a trimer that constitutes the distal part of the tail fibers and attaches to *Listeria* cells before sheath contraction. Therefore, it is likely that WD0577 plays a role in host binding and may form part of the tail fiber.

Finally, **WD0576** is predicted to act as a *Caudoviricetes* tail fiber assembly chaperone similar to GpK (HHpred, 92.7% probability, E-value 4.3). In lambda phages, GpK forms a scaffold at the tip of the tail tube where tail fibers or receptor binding proteins self-assemble [25]. Tail fiber assembly chaperones are typically located downstream from tail fiber proteins [26]. Therefore, based on gene synteny and homology to *Listeria* phage A511, WD0580/WD0579 and WD0577 are tail fiber candidates whereas WD0576 and WD0578 are chaperone candidates of phage WO's tail fiber module.
