## Supplementary figures and images for "The most widespread phage in animals: Genomics and taxonomic classification of Phage WO"

### S1 Fig

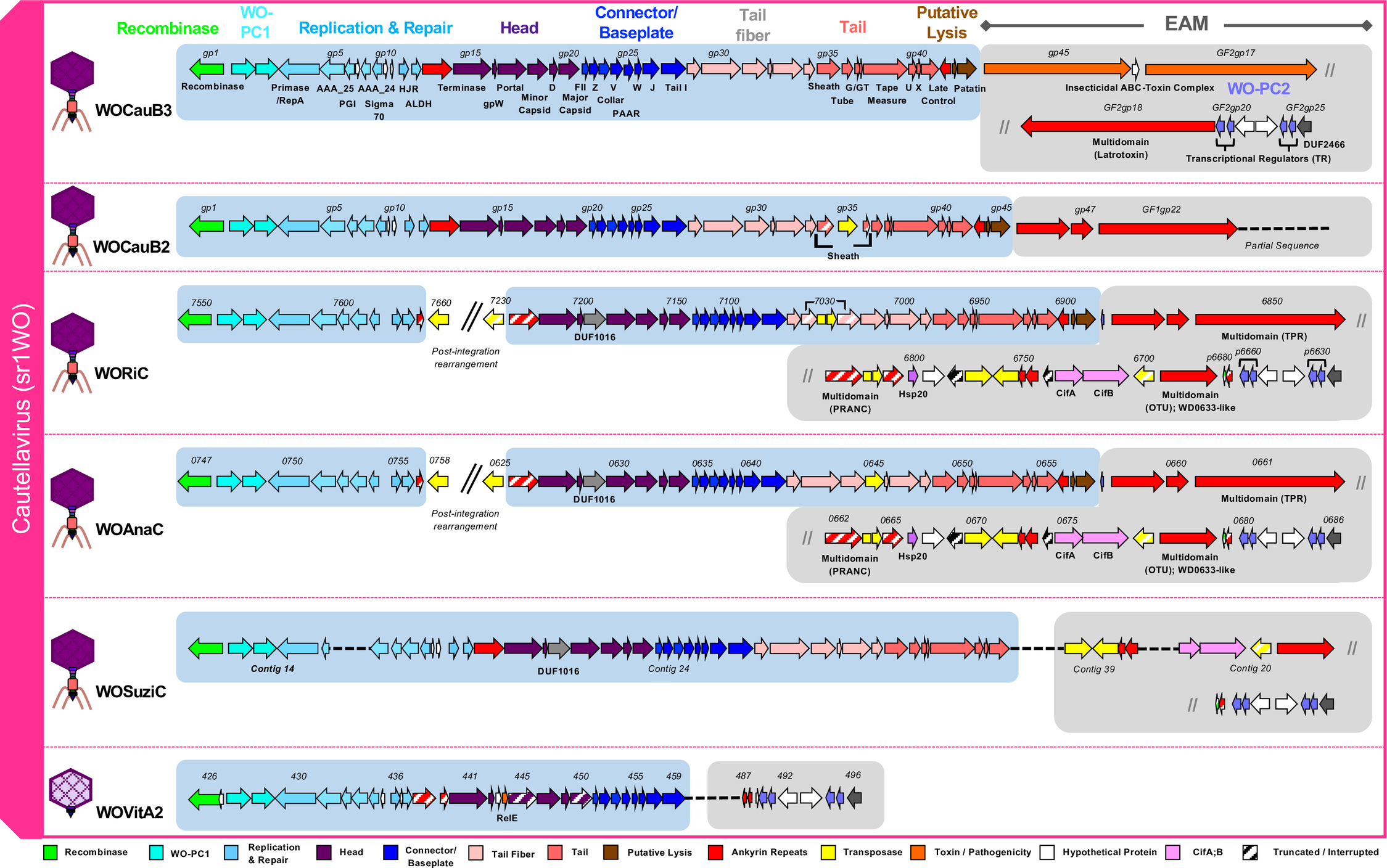

### S2 Fig

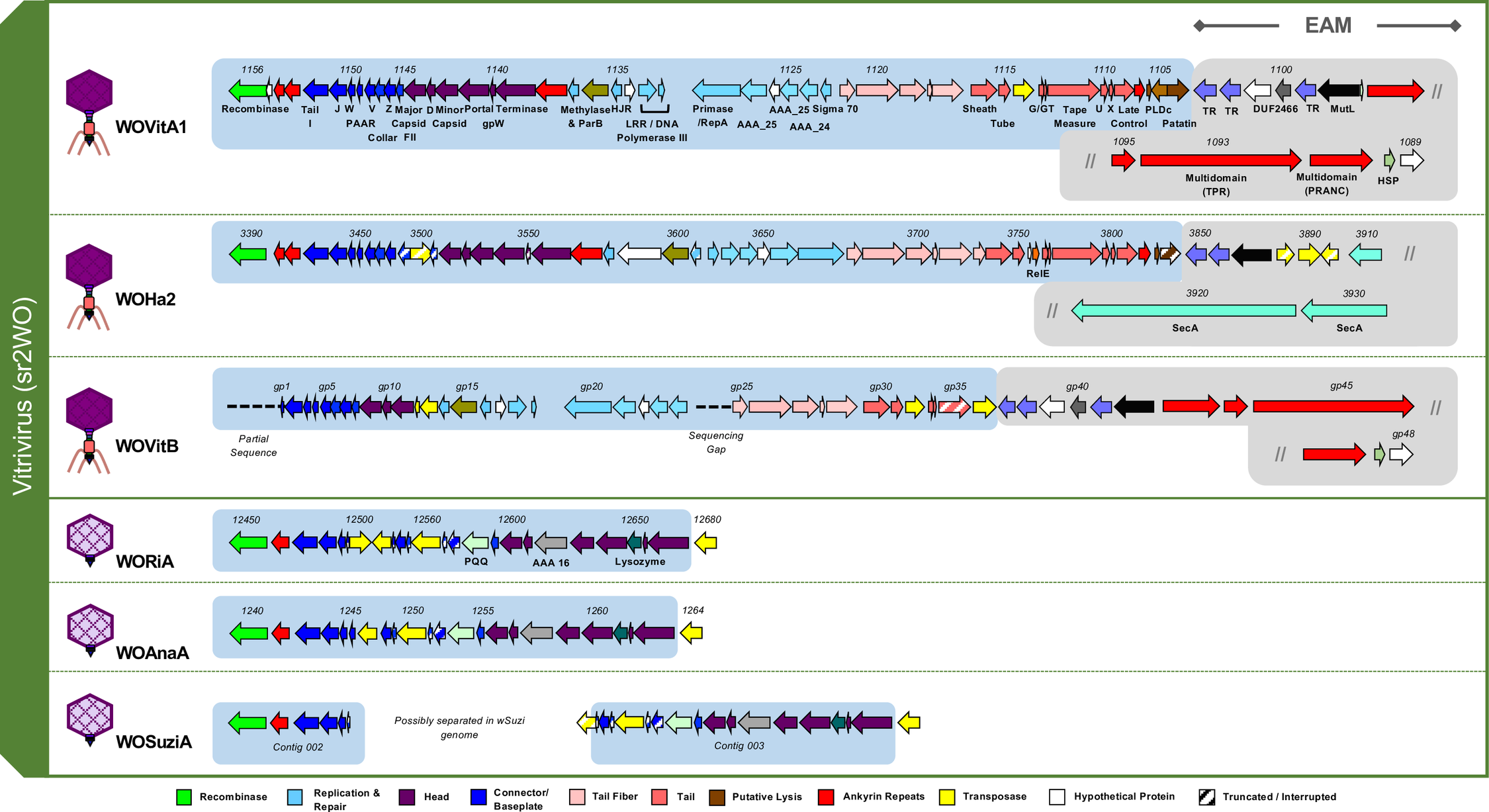

### S3 Fig

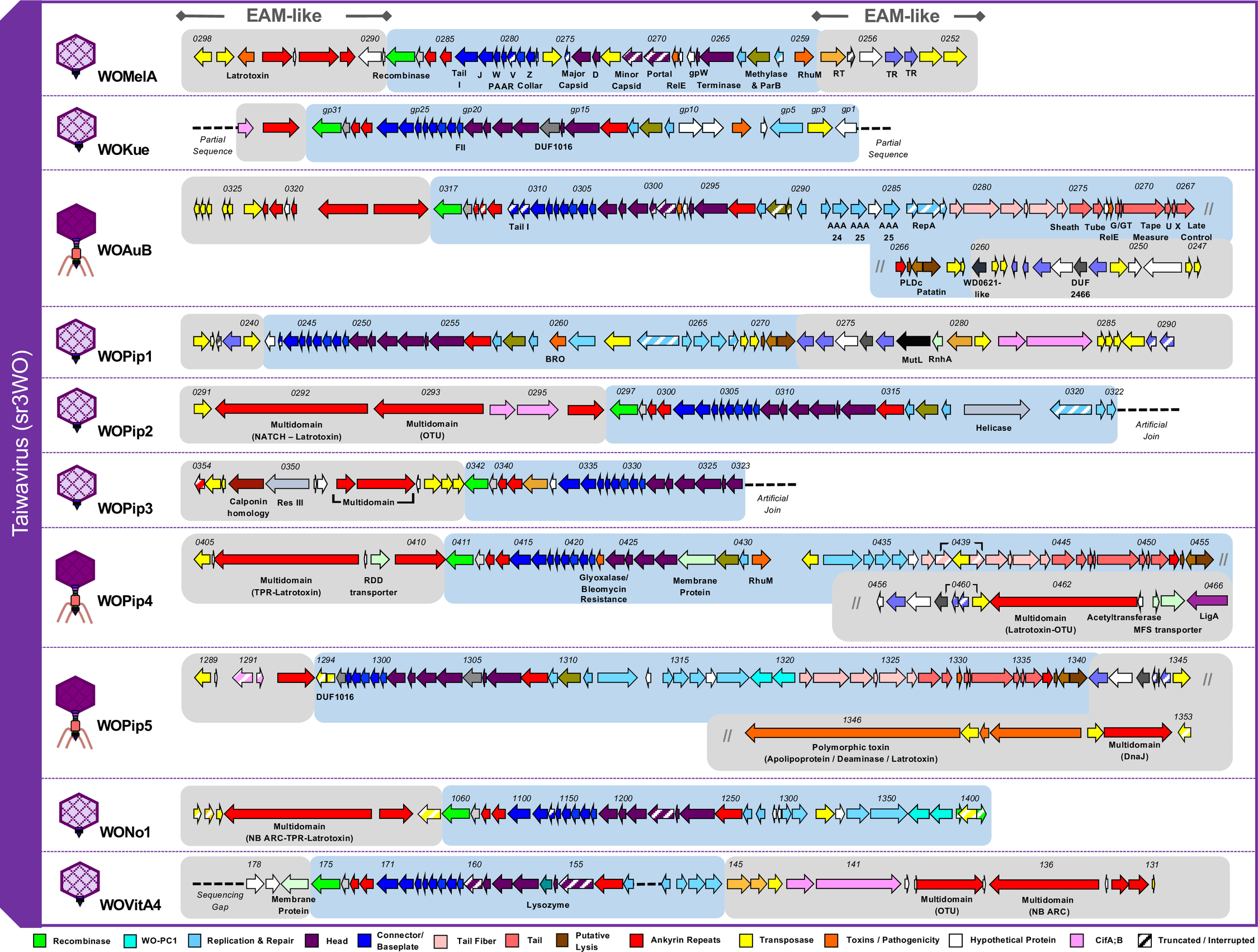

### S4 Fig

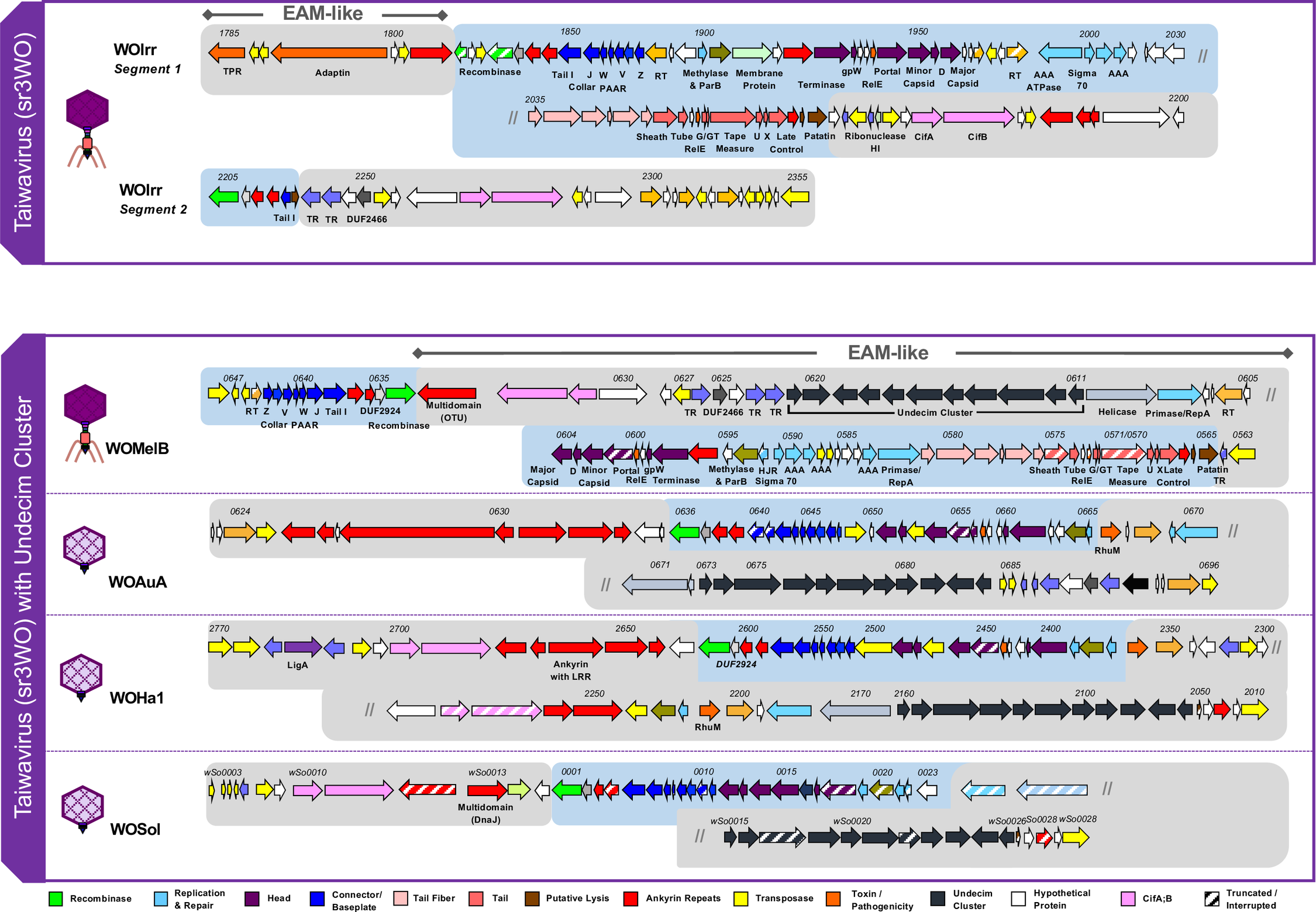

### S5 Fig

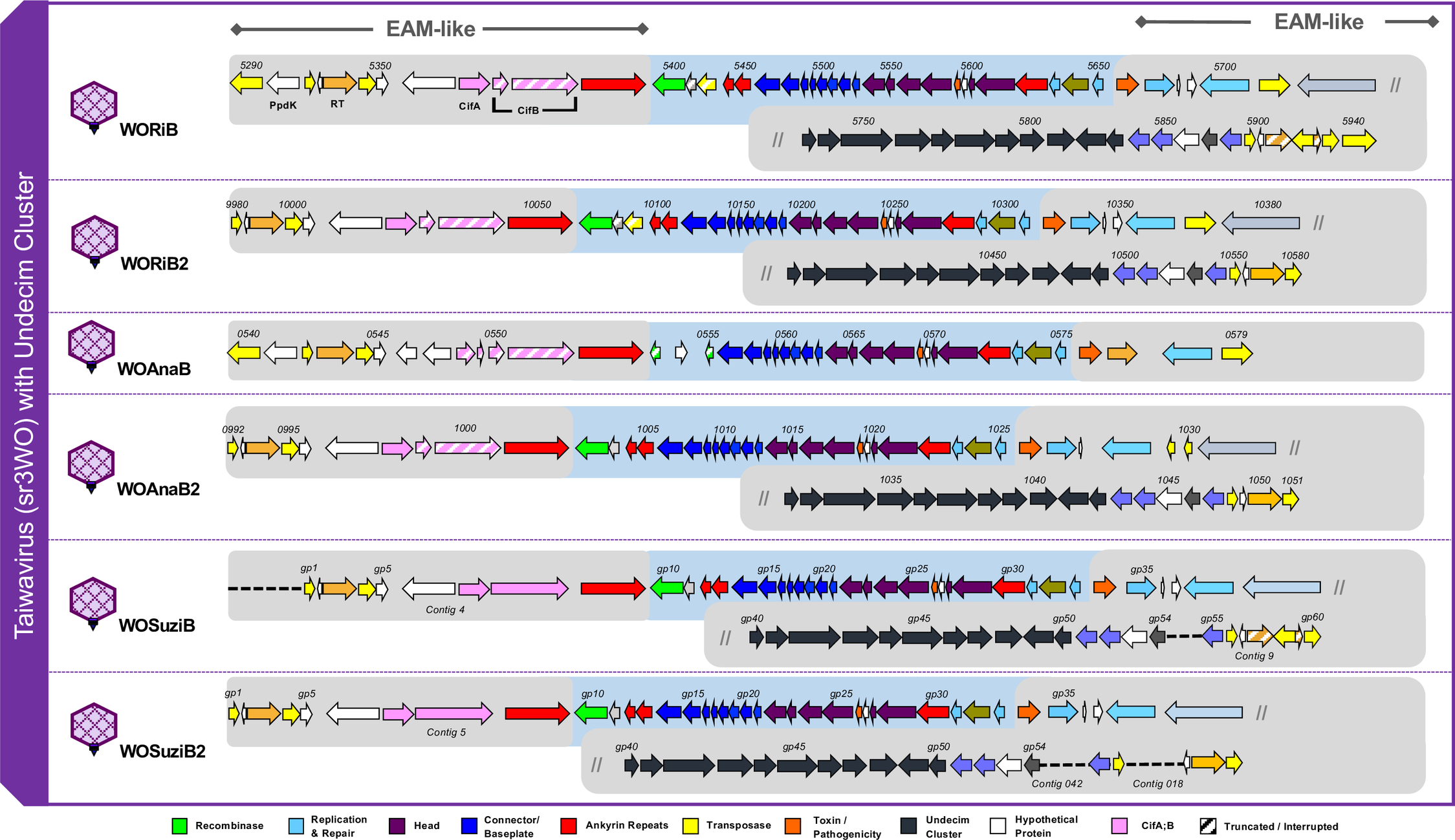

### S6 Fig

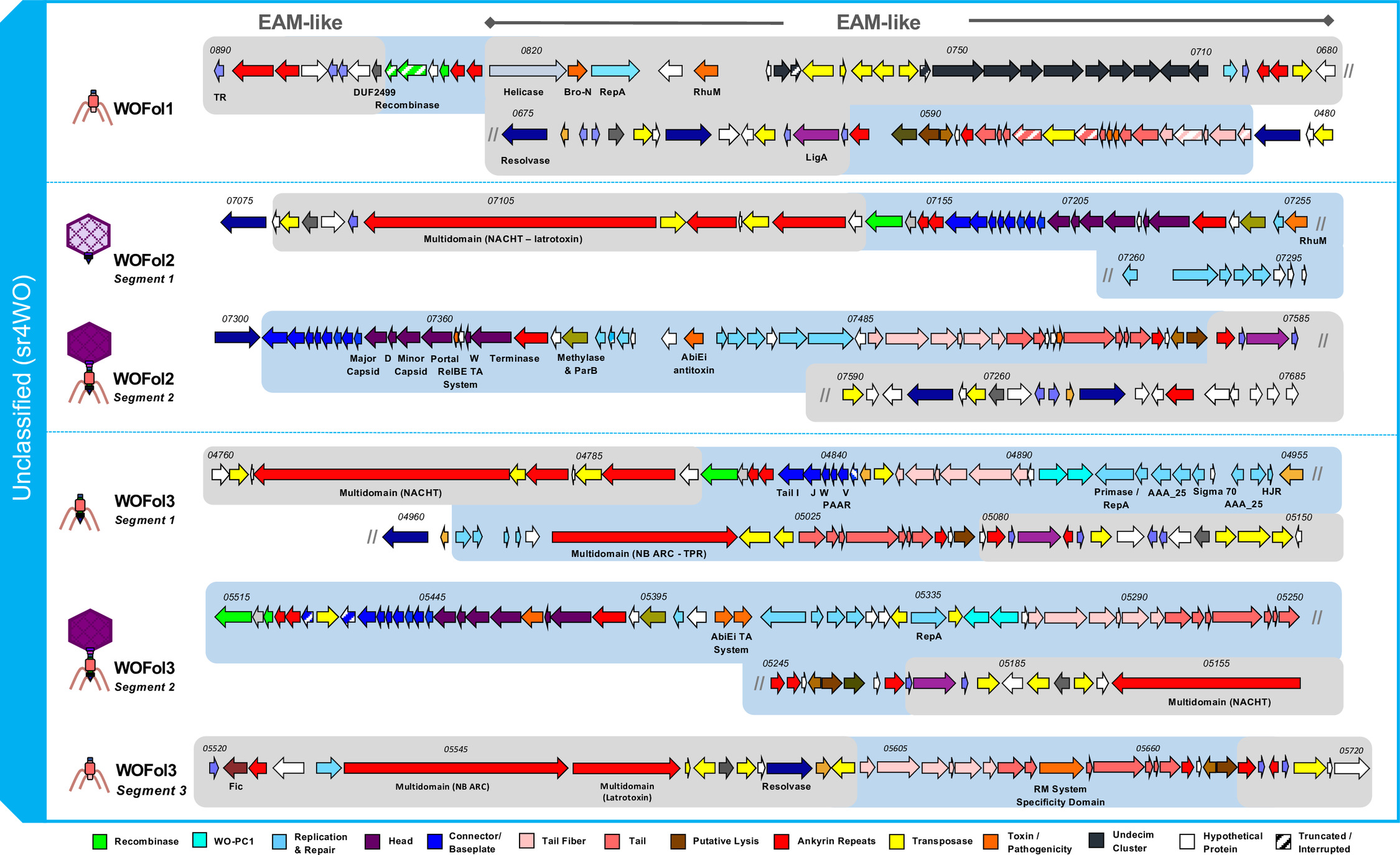

### S7 Fig

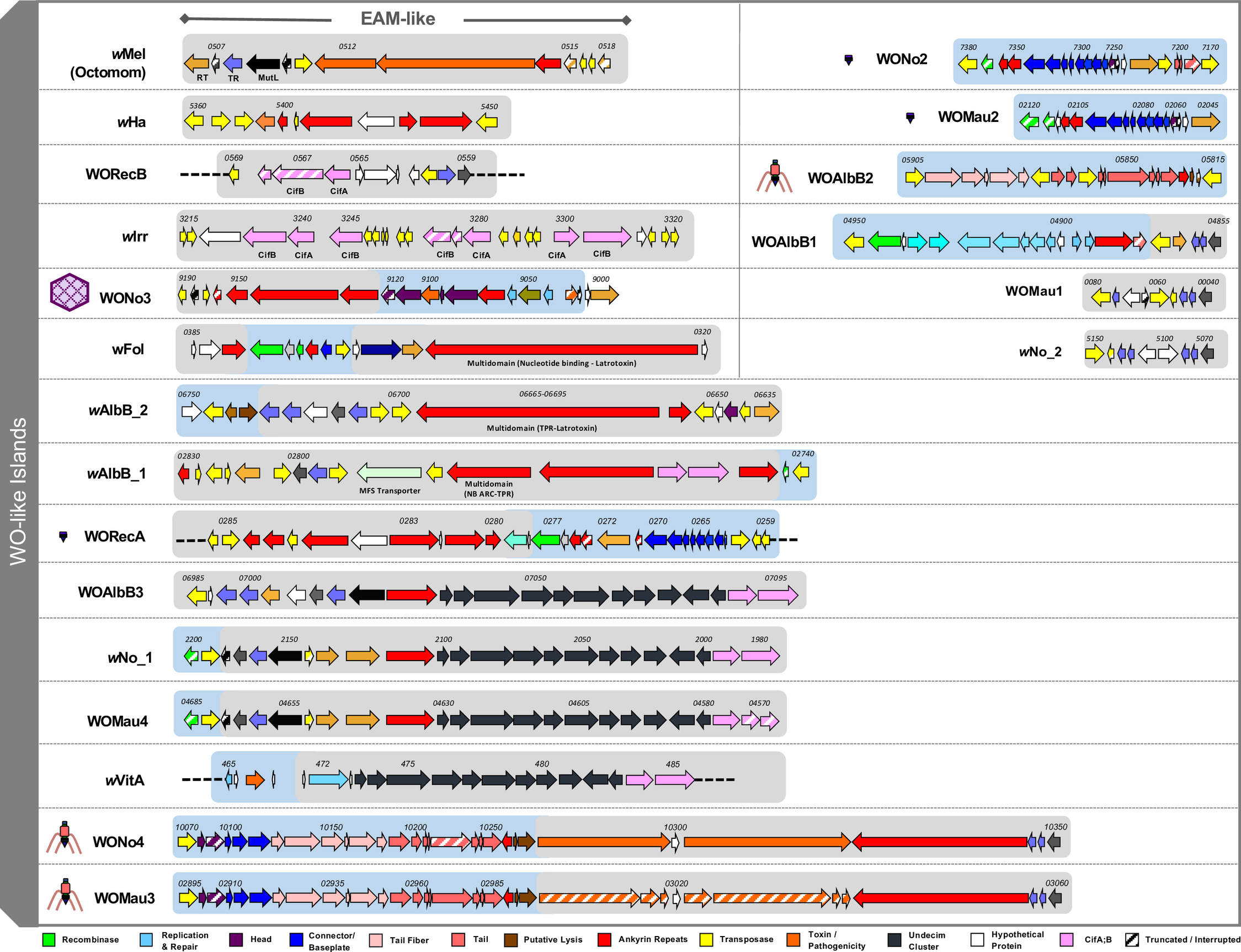

### S8 Fig

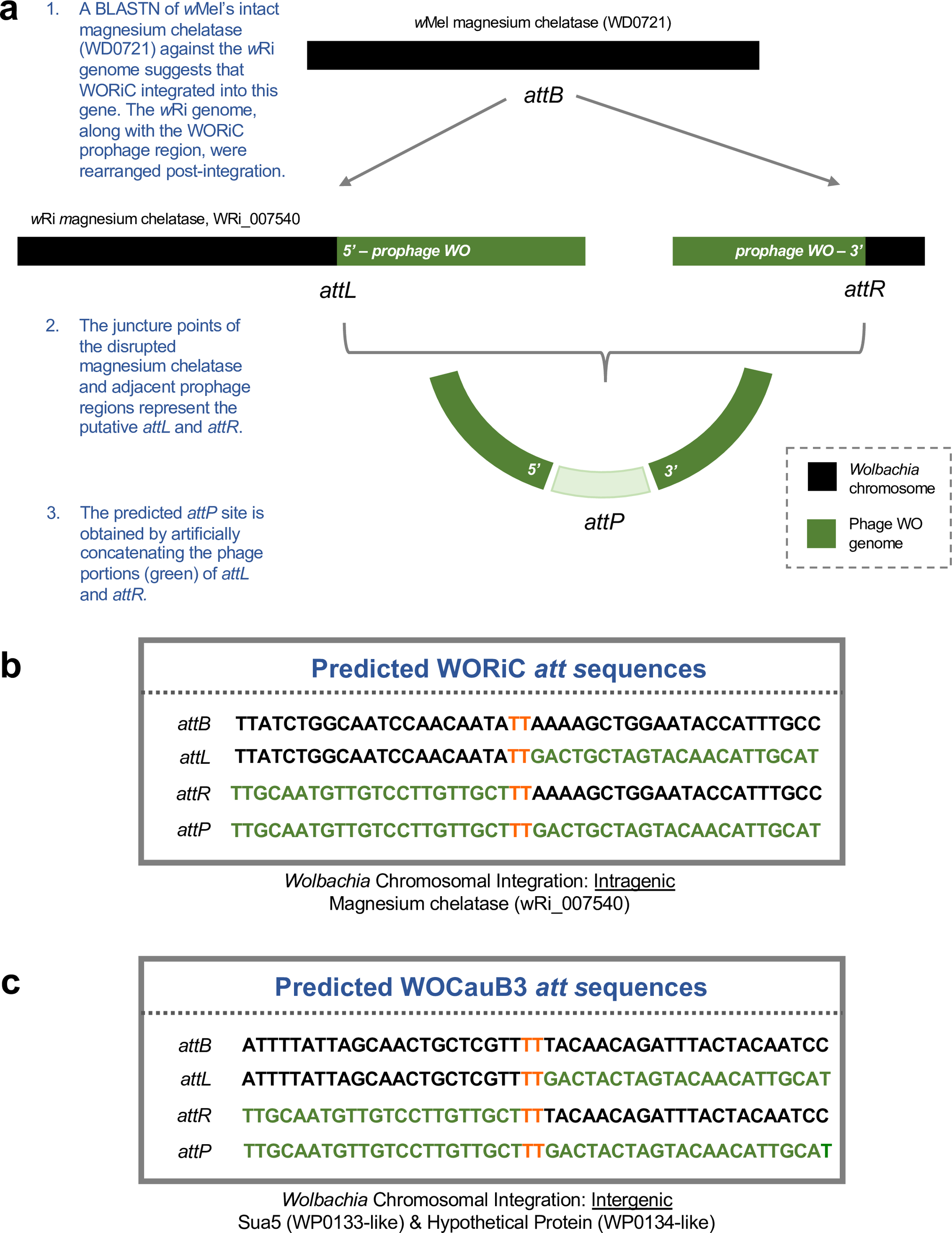

### S9 Fig

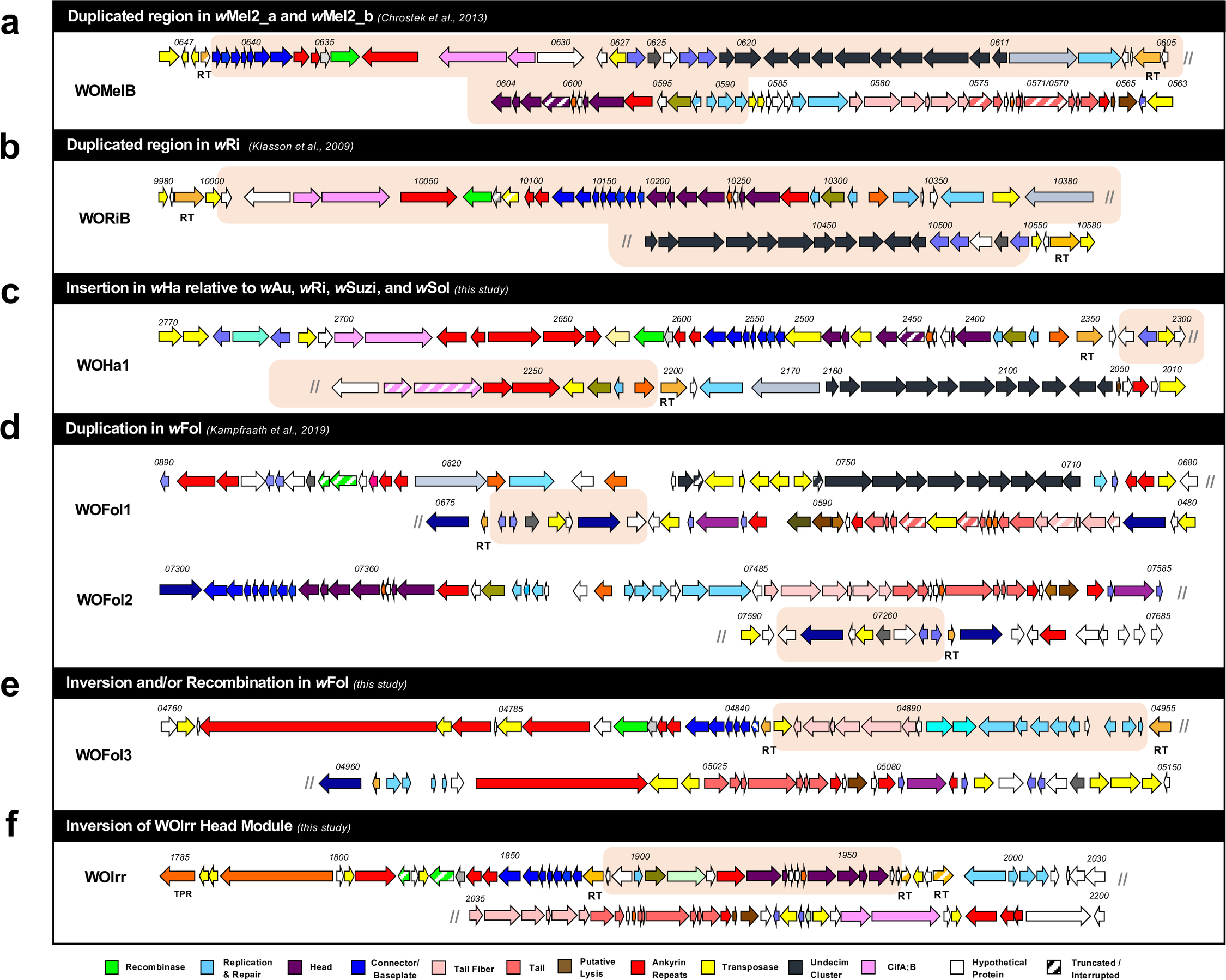

### S10 Fig

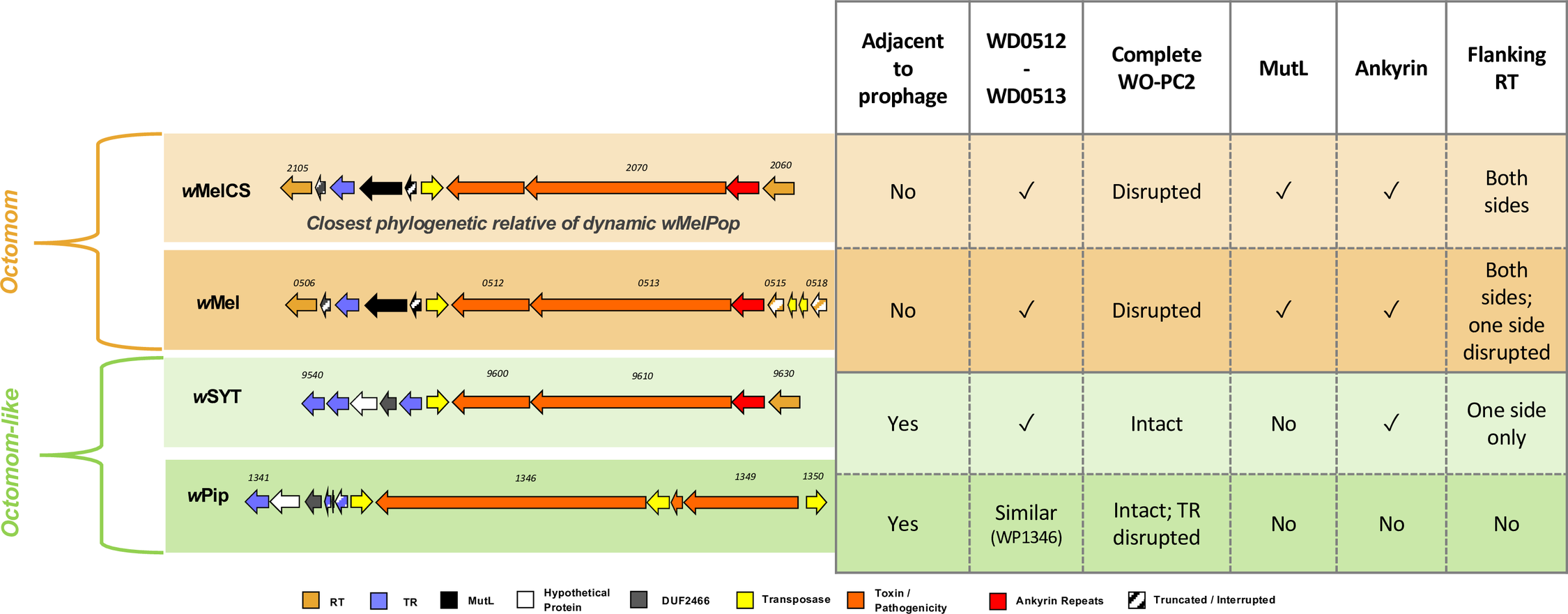

### S11 Fig

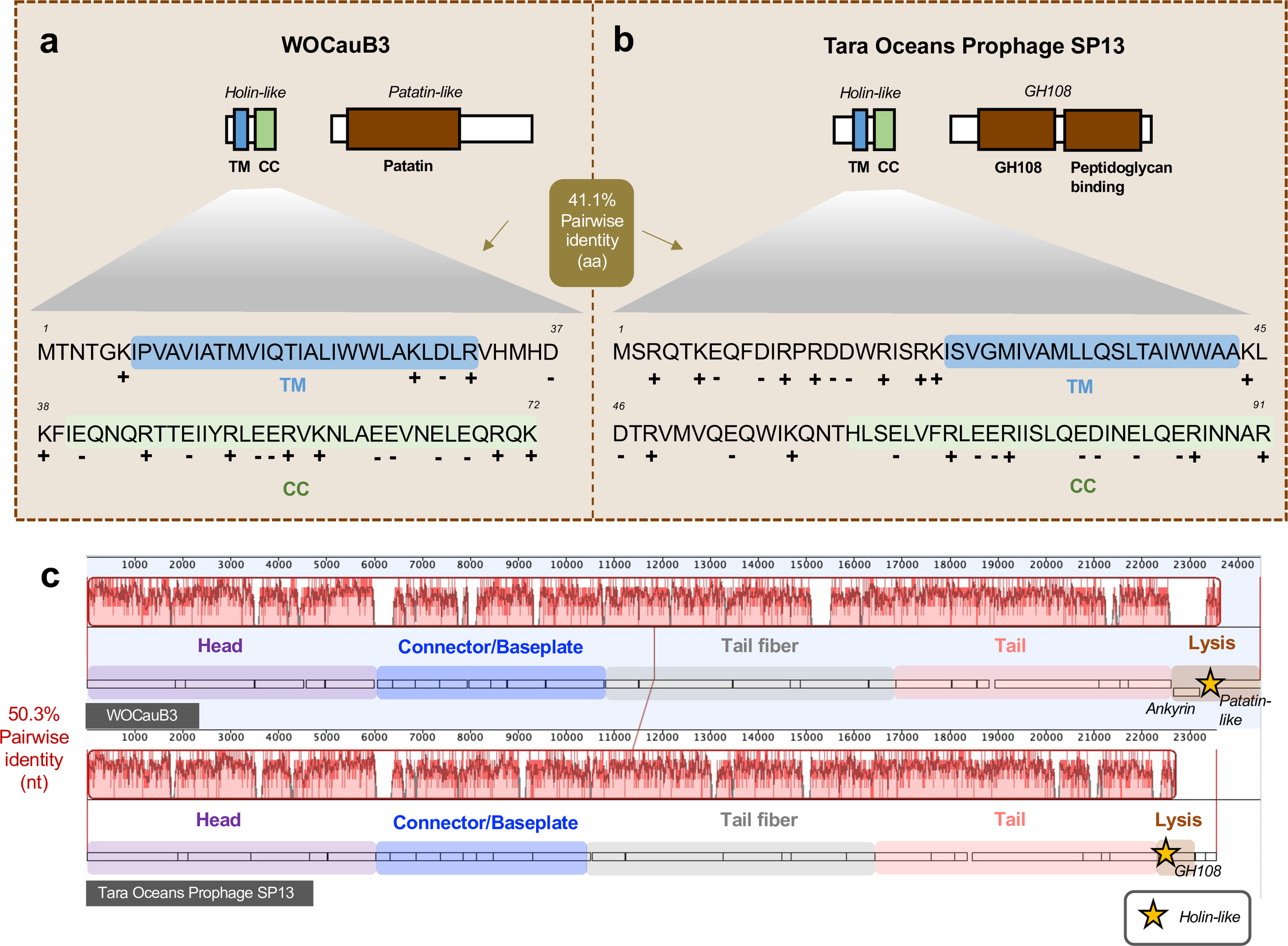

### S12 Fig

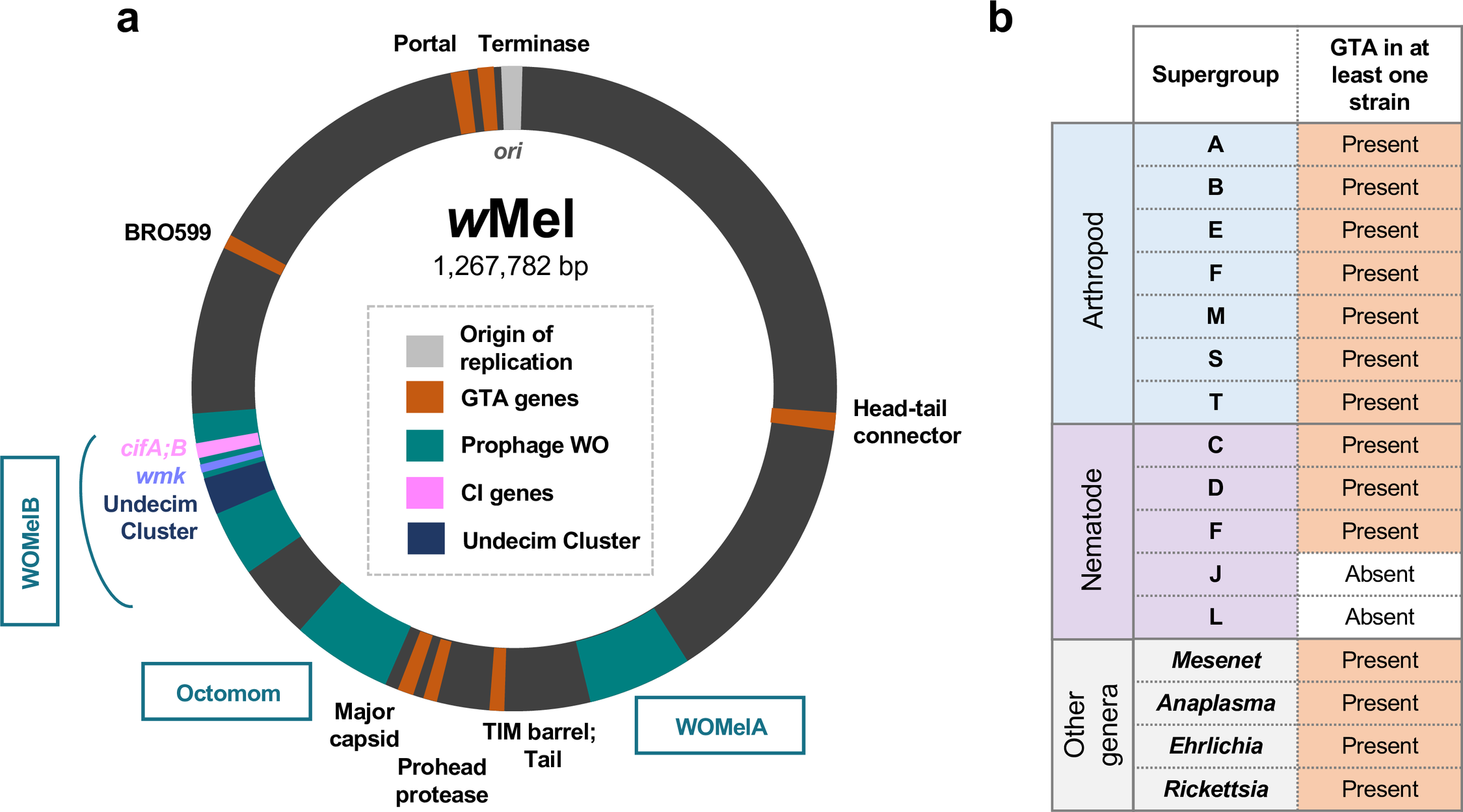

### S13 Fig

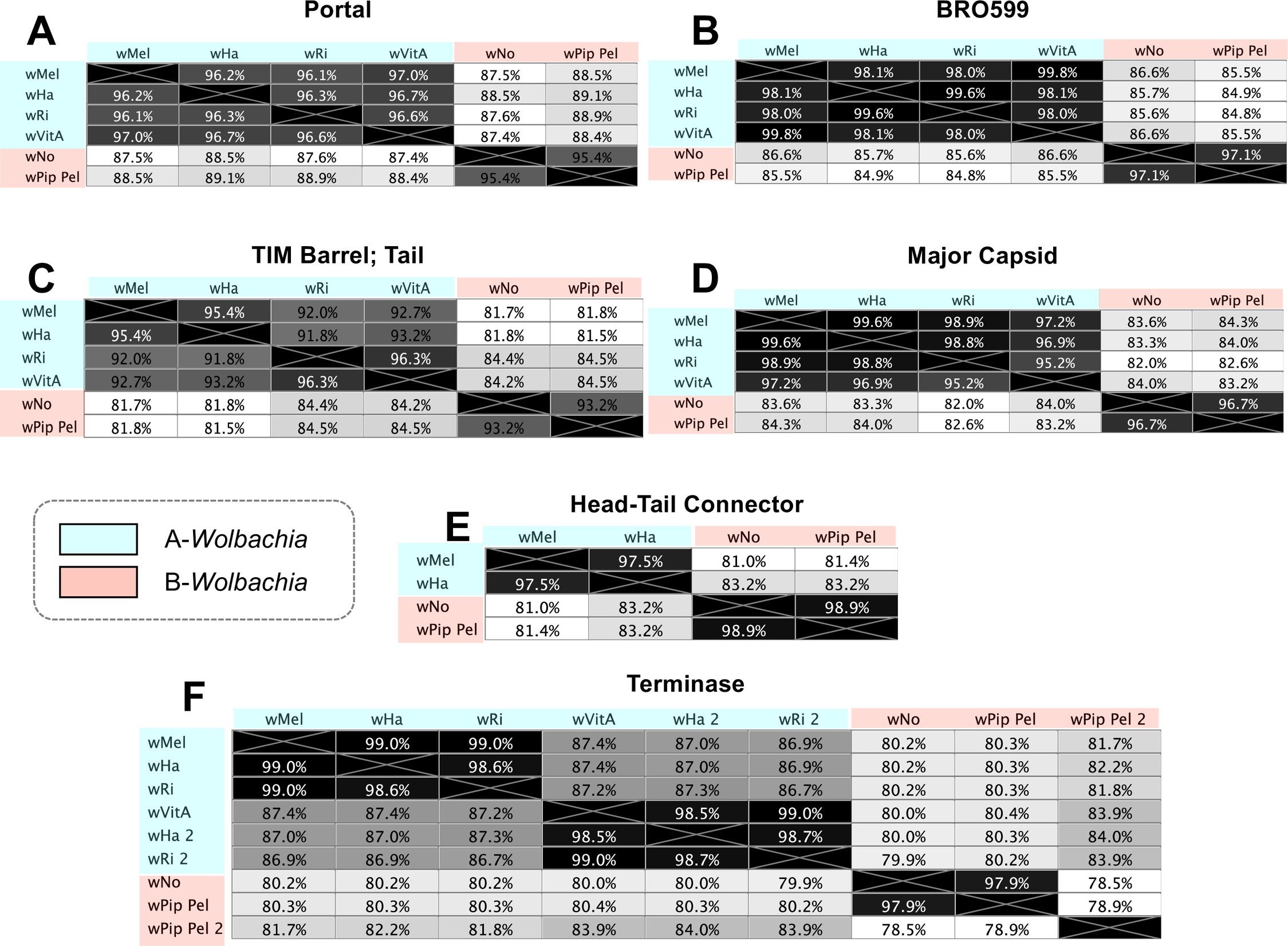
